# *In-situ* structure of the flagellar export apparatus in *Borrelia burgdorferi*

**DOI:** 10.64898/2026.05.13.724734

**Authors:** Jian Yue, Akarsh Manne, Wangbiao Guo, Huaxin Yu, Jiaqi Wang, Md Khalesur Rahman, Kathryn Lees, Hui Xu, Jack M. Botting, Md A. Motaleb, Jun Liu

## Abstract

The transmembrane export apparatus is a conserved core of bacterial type III secretion systems, shared by the flagellum and injectisome. Powered by the proton-motive force (PMF), this machinery translocates protein substrates across the bacterial envelope to support motility and virulence. However, its assembly and operation within native membranes remain poorly understood. Here, using *in-situ* single-particle cryo-electron microscopy, we resolve near-atomic-resolution structures of the flagellar export apparatus within the periplasmic flagellar motor of the Lyme disease spirochete *Borrelia burgdorferi*. Structural and genetic analyses suggest that coordinated, stepwise assembly of the flagellar export apparatus and MS-ring drives local deformation of the cytoplasmic membrane through extensive protein-lipid-protein interactions, constructing a funnel-shaped conduit optimal for substrate translocation. In addition, a conserved hydrophilic pathway lies within core export protein FlhA, with key acidic residues essential for export activity and supporting PMF-driven substrate translocation. Together, these findings establish a mechanistic framework for export apparatus assembly and energy transduction, defining core principles of type III secretion.

## MAIN

Bacterial flagella and injectisomes are sophisticated, self-assembling nanomachines whose origins trace to an ancient flagellum in the last bacterial common ancestor^1,2^. The flagellum has evolved to play centrals in motility, host-pathogen interaction, and pathogenesis across diverse bacterial species, while injectisomes have specialized to directly translocate effector proteins into host cells, promoting infection^3,4^. A central component shared by both nanomachines is the type III secretion system (T3SS)^5-7^, which is powered by proton-motive force (PMF) and ATP hydrolysis^8,9^, enabling substrate unfolding and translocation across the membrane. Unlike the general Sec secretion system, which translocates ∼40 amino acids per second^10^, the T3SS is highly efficient, exporting up to 10,000 amino acids per second during initial flagellar growth, with the export rate decreasing rapidly as filament length increases in *Salmonella enterica*^11,12^. However, the molecular mechanisms underlying type III secretion remain unclear^13^.

The flagellar T3SS consists of a transmembrane export apparatus formed by five membrane proteins (FlhA, FlhB, FliP, FliQ, and FliR) and a cytoplasmic ATPase complex (FliH, FliI, and FliJ)^14,15^. Structural studies have shown that FliP, FliQ, and FliR assemble into a helical export gate that forms the central protein-conducting channel of the system^16,17^, while FlhB associates with this complex and contributes to substrate specificity switching during flagellar assembly^18,19^. FlhA is the largest component of the export apparatus and forms a nonamer that interacts with substrates, chaperones, and the ATPase complex^20-25^. Genetic and biochemical studies further suggest that FlhA plays a central role in coupling PMF to protein export^26-29^. However, recent export apparatus structures from purified basal bodies of *S. enterica*^30,31^ do not fully capture its organization and function within the native membrane environment. In particular, how FlhA is spatially arranged relative to the FliPQR–FlhB export gate and how this organization supports PMF-driven protein export remain unresolved^13^.

Here, we combine *in-situ* single-particle cryo-electron microscopy (cryo-EM) and cryo-electron tomography (cryo-ET) with molecular genetics and modeling to reconstruct the intact transmembrane export apparatus within the periplasmic flagellar motor of *Borrelia burgdorferi*, the causative agent of Lyme disease. Our *in-situ* study reveals a near-atomic-resolution structure of the export apparatus and the associated MS-ring in intact bacteria and supports a model in which coordinated, sequential assembly of both drives membrane deformation, generating a funnel-shaped conduit through protein-lipid-protein interactions. Structural and mutational analyses further provide evidence that a conserved hydrophilic pathway within FlhA is essential for PMF-driven protein secretion. Together, our results establish that coordinated, sequential assembly of the export apparatus and MS-ring plays a crucial role in early flagellar biogenesis and enables efficient type III secretion.

## RESULTS

### *In-situ* structure of the export apparatus and MS-ring

To visualize the flagellar export apparatus in its native membrane context, we first determined the *in-situ* structure of the flagellar motor from wild-type *B. burgdorferi* cells, using cryo-ET and subtomogram averaging (Fig. 1a). The export apparatus and surrounding MS-ring are embedded within a highly curved cytoplasmic membrane, an architecture not previously characterized in bacterial flagellar systems^32,33^. The membrane underneath the MS-ring bends sharply toward the cytoplasmic side, forming a pronounced hemispherical deformation with a depth of 10 nm (Fig. 1a). Focused refinements further resolve a distinct membrane curvature in which the spacing between two leaflets of the cytoplasmic membrane ranges from 3.3nm at the periphery to 2.5nm adjacent to the FliPQR–FlhB complex (Fig. 1b,c; Extended Data Fig. 1a,b). These findings suggest that the flagellar export apparatus and surrounding MS-ring function as a membrane-sculpting complex.

**Fig. 1:**
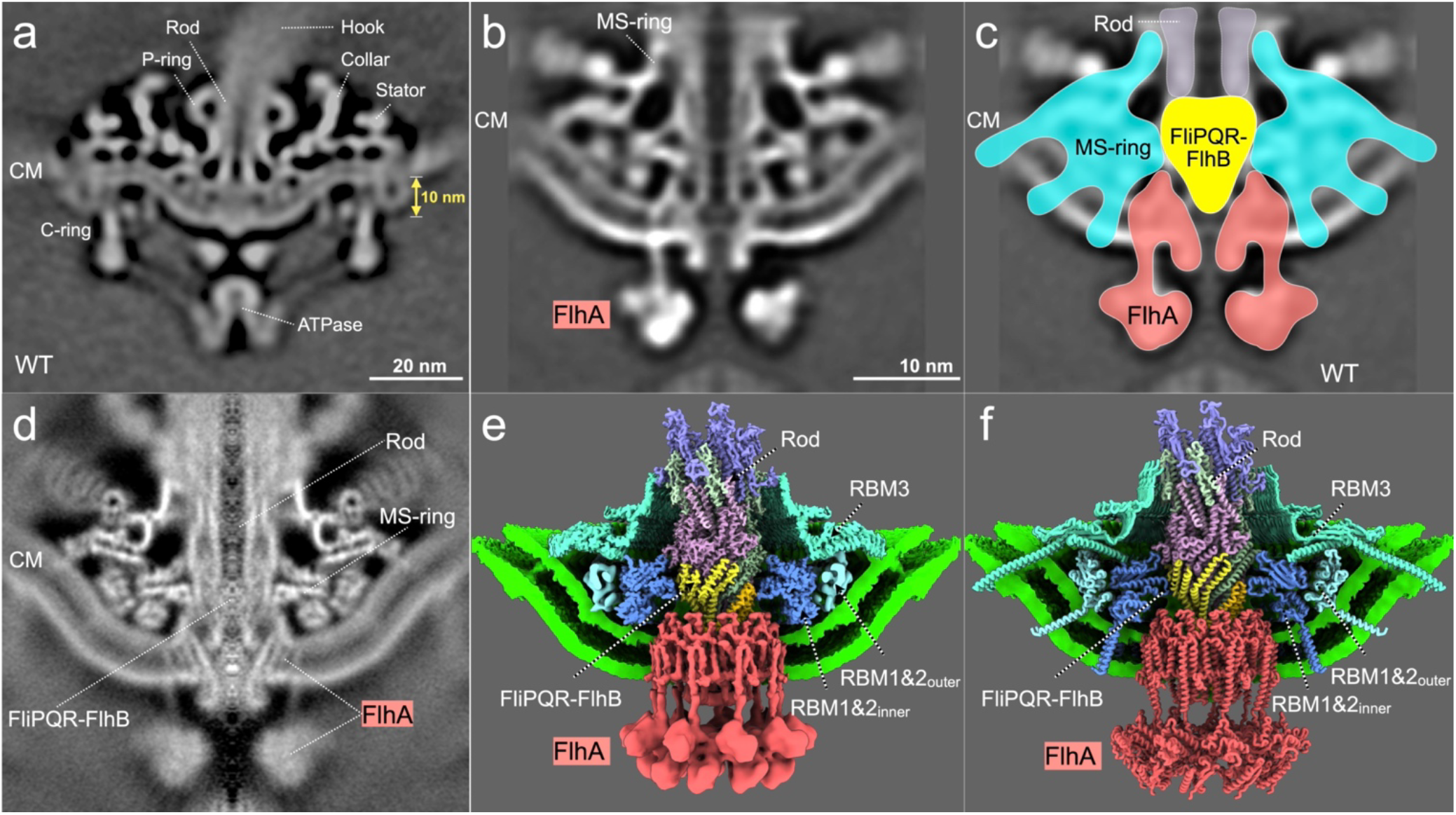
*In-situ* architecture of the flagellar export apparatus within a highly curved cytoplasmic membrane. (a) Central slice of the flagellar motor from wild-type (WT) cells determined by cryo-ET and subtomogram averaging, with major components labeled, including the C-ring, stator, collar, ATPase complex, and MS-ring. CM, cytoplasmic membrane. (b) Focused refinement of the export apparatus and MS-ring. (c) Schematic illustrating the organization of the export apparatus within the MS-ring (FliF). FliPQR–FlhB forms a cone-shaped core, with FlhA adopting a seahorse-like conformation beneath. The cytoplasmic membrane is highly curved. (d) Central section of the export apparatus and MS-ring resolved by *in-situ* single-particle cryo-EM. (e) Composite high-resolution structure. RBM3 (cyan) exhibits C41 symmetry (3.4 Å), whereas the FliPQR–FlhB gate, proximal rod, and FliF_RBM1–RBM2_ inner ring were resolved at 3.8 Å without symmetry (C1). The FliF_RBM1–RBM2_ outer ring shows C18 symmetry (6.5 Å). FlhA_TMD_ nonamer was resolved by cryo-EM at 4.3 Å resolution, whereas FlhA_CTD_ was resolved by cryo-ET at 16 Å resolution. (f) Overall structure of the export apparatus and MS-ring. The MS-ring embedded in the curved cytoplasmic membrane is shown in a central cross-section.

To resolve the molecular details of this membrane-sculpting complex, we implemented an *in-situ* single-particle cryo-EM strategy^34^ to analyze 43,970 flagellar motor particles extracted from 26,472 micrographs (Fig. 1d,e). Through focused classification and refinement, we resolved nearly the entire MS-ring and the export apparatus at near-atomic resolution. The MS-ring is assembled from several FliF domains with distinct symmetries (Fig. 1e; Extended Data Figs. 2, 3, 4). The FliF ring-building motifs FliF_RBM1_ (residues 51-110) and FliF_RBM2_ (residues 111-225) organize into two concentric rings: an inner ring with C23 symmetry resolved at 3.5 Å and a more flexible outer ring with C18 symmetry resolved at 6.5 Å (Fig. 1e; Extended Data Figs. 2, 3). FliF_RBM3_ (residues 226-450) forms the third ring with C41 symmetry resolved at 3.4 Å (Fig. 1e; Extended Data Figs. 2, 3). Each FliF subunit contains two transmembrane helices, FliF_TMI_ (residues 25-45) and FliF_TMII_ (residues 473-493) (Extended Data Fig. 4). These helices exhibit well-defined densities that traverse the curved lipid bilayer and connect to the FliF_RBM1–RBM2_ and FliF_RBM3_ rings, respectively (Fig. 1e; Extended Data Fig. 4g-i; Movie S1). In total, the MS-ring comprises 82 transmembrane helices arranged in three concentric layers: 23 FliF_TMI_ helices from the inner FliF_RBM1–RBM2_ ring, 18 FliF_TMI_ helices from the outer FliF_RBM1–RBM2_ ring, and 41 FliF_TMII_ helices forming the outermost layer (Fig. 1f; Fig. 2a,b; Extended Data Fig. 4j,k). Together, these helices with hydrophobic surfaces enable formation of the curved membrane (Fig. 2c). Notably, the resulting *B. burgdorferi* MS-ring is substantially larger and more complex than that of *S. enterica* (Extended Data Fig. 4l,m).

**Fig. 2:**
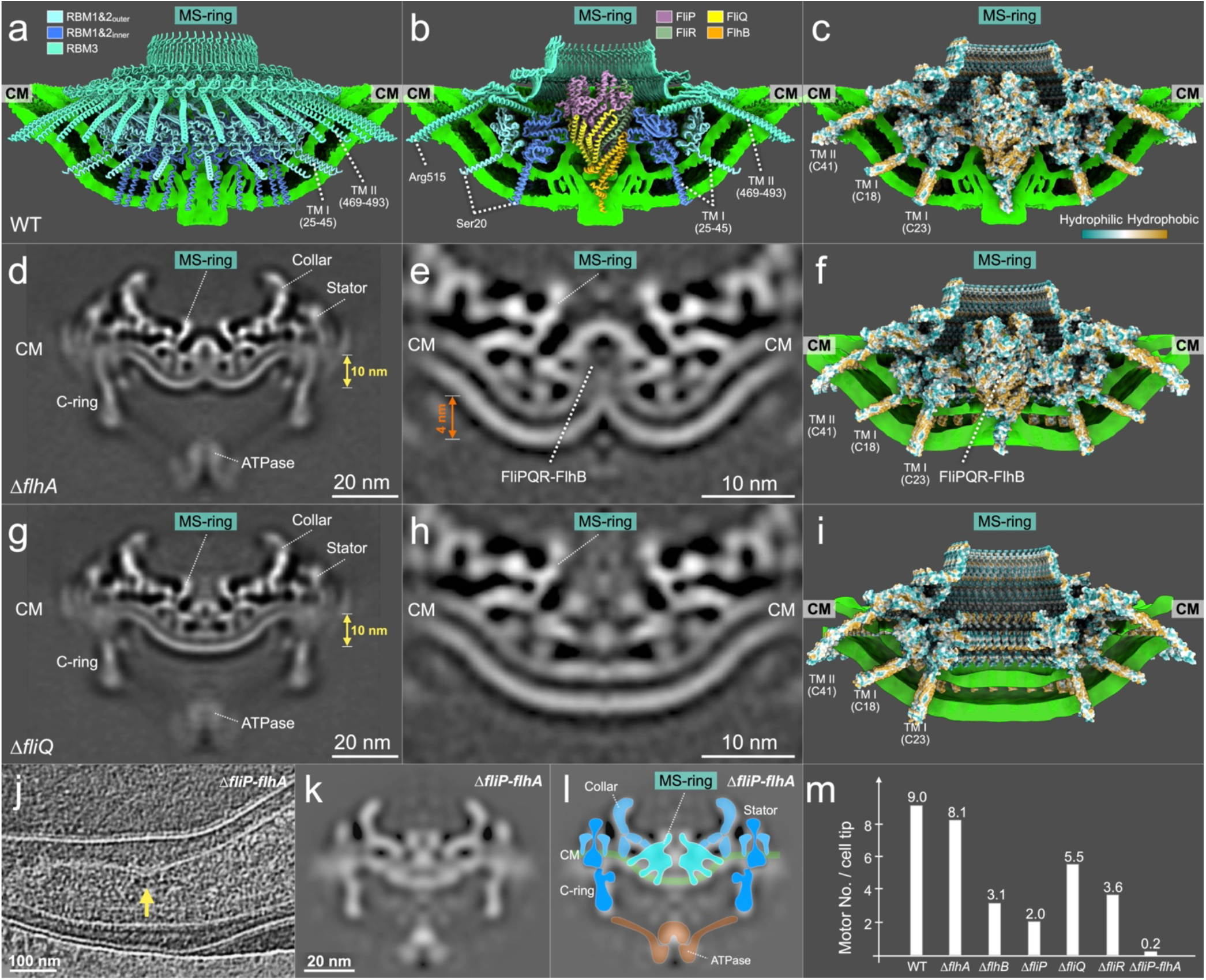
Coordinated assembly of the MS-ring and export apparatus drives membrane deformation. (a) Model of the MS-ring with transmembrane helices extended based on *in-situ* density and AlphaFold3 predictions. (b) Central cross-section showing the MS-ring surrounding the membrane-embedded FliPQR–FlhB export gate. (c) Central cross-section showing the MS-ring and the export gate with hydrophobic surfaces aligned to the membrane curvature. (d-f) In the Δ*flhA* mutant, the MS-ring remains assembled but is embedded within a pronounced W-shaped membrane, with hydrophobic surfaces aligned to membrane curvature and the export gate elevated by 4nm. (g-i) In the Δ*fliQ* mutant, the MS-ring is embedded within a U-shaped membrane, contrasting with the W-shaped curvature observed in wild-type and Δ*flhA* cells, demonstrating that MS-ring and export-apparatus assembly drive distinct modes of membrane deformation. (j) Cryo-ET slice from a cell tip of the Δ*fliP–flhA* mutant. (k, l) Central section and corresponding model of the motor structure in the Δ*fliP–flhA* mutant. (m) Quantification shows motor numbers per cell tip in wild type and six mutants.

The FliPQR–FlhB export gate in complex with the proximal rod (FliE, FlgB, and FlgC)^35^ and FliF_RBM1–RBM2_ inner ring were determined at 3.8 Å (Fig. 1e; Extended Data Figs. 2, 3). In parallel, the FlhA_TMD_ nonamer structure with C9 symmetry was resolved at 4.3 Å (Fig. 1e; Extended Data Figs. 2, 3). Although the FlhA_CTD_ nonamer was not well resolved by *in-situ* single-particle cryo-EM, likely due to conformational flexibility or interference from the cellular background, it was visualized at ∼16 Å resolution by cryo-ET and subtomogram averaging (Fig. 1a-c). Together, these datasets enabled us to construct a composite model of the intact export apparatus within the MS-ring (Fig. 1f; Extended Data Fig. 5; Movie S2). This composite model includes 6 FliE, 5 FlgB, and 6 FlgC subunits in the proximal rod; 5 FliP, 1 FliR, 4 FliQ, 1 FlhB, and 9 FlhA subunits in the export apparatus; and 41 FliF subunits in the MS-ring, providing a structural basis for understanding the assembly and function of the export apparatus in the context of the locally curved cytoplasmic membrane.

### Assembly of the MS-ring and export apparatus drives profound membrane deformation

To better understand how the MS-ring and export apparatus are embedded in the curved membrane, we determined the motor structures in the absence of each component of the export apparatus using cryo-ET and subtomogram averaging (Fig. 2d,e,g,h; Extended Data Fig. 6). Different from the wild-type motor, the Δ*flhA* motor lacks the rod, hook, and filament (Fig. 2d-f). The cytoplasmic membrane in the Δ*flhA* motor adopts a pronounced W-shaped cross-section with the spacing between leaflets ranging from 3.3nm at the periphery to 2.5nm adjacent to the FliPQR–FlhB complex (Fig. 2d-f; Extended Data Fig. 1c,d). In this configuration, the hydrophobic surface of the FliPQR–FlhB complex is positioned ∼4 nm above the lowest level of the cytoplasmic membrane (Fig. 2e,f). This local membrane deformation accommodates the elevated FliPQR–FlhB complex as well as likely enables stable assembly of the FlhA nonamer. Consistent with this notion, FlhA fails to assemble in the Δ*fliQ* motor, in which the MS-ring is embedded in a U-shaped cytoplasmic membrane with the spacing between leaflets ranging from 3.2nm to 3.5nm (Fig. 2g-i; Extended Data Fig.1e,f). Notably, the MS-ring remains structurally indistinguishable across wild-type, Δ*flhA*, Δ*fliQ* motors (Fig. 2) as well as in Δ*fliP* and Δ*fliR* mutants (Extended Data Fig. 6), suggesting that MS-ring assembly drives local membrane deformation, while interactions between the MS-ring and the export gate elevate the intervening membrane, collectively generating the unique export apparatus architecture embedded in the curved membrane.

### The transmembrane export apparatus functions as a template for efficient MS-ring assembly

To dissect how the export apparatus and MS-ring coordinate their assemblies in driving membrane deformation, we constructed a quintuple mutant (Δ*fliP-flhA*), which eliminates the entire transmembrane export apparatus but does not affect the expression levels of FliF (Fig. 2j,k,l; Extended Data Fig. 7). Strikingly, compared to wild type and single-gene deletion mutants, the motors in the quintuple mutant are less visible. Only 54 motors were identified from 342 reconstructions of the quintuple mutant cell tips (Fig. 2m). In total, the number of motors is reduced by ∼98% compared with wild type, a far more severe defect than that produced by any single-gene deletion (Fig. 2m). Our *in-situ* motor structure clearly shows that the rod, hook, and filament fail to assemble in the mutant (Fig. 2k,l), indicating that assembly of the MS-ring is not strictly dependent on the export apparatus but becomes inefficient or unfavorable in its absence (Fig. 2j-m). Together, these findings support a model in which the export apparatus plays dual roles: it is indispensable for T3SS-mediated assembly of the rod, hook, and filament, and it also functions as a transmembrane nucleation template that accelerates and stabilizes MS-ring assembly, thereby ensuring efficient initiation of flagellar biogenesis (Fig. 2m).

### *In-situ* structure of the FlhA nonamer in a locally curved membrane

FlhA assembles as a nonamer embedded within the locally deformed, W-shaped membrane surrounding the export gate (Fig. 1d-f; Fig. 3a). Each FlhA subunit comprises a FlhA_TMD_ and FlhA_CTD_ connected by a linker helix. The FlhA_TMD_ contains eight helices (H1-H8) organized into a unique architecture: H1-H3 and H7-H8 occupy a lower position in the bilayer, while H4-H6 rise by up to ∼30 Å, generating a stepped, ramp-like hydrophobic surface across the membrane (Fig. 3c,d). Structural comparisons by Foldseek^36^ and DALI^37^ indicate that this architecture is conserved across T3SSs (Extended Data Fig. 8).

**Fig. 3:**
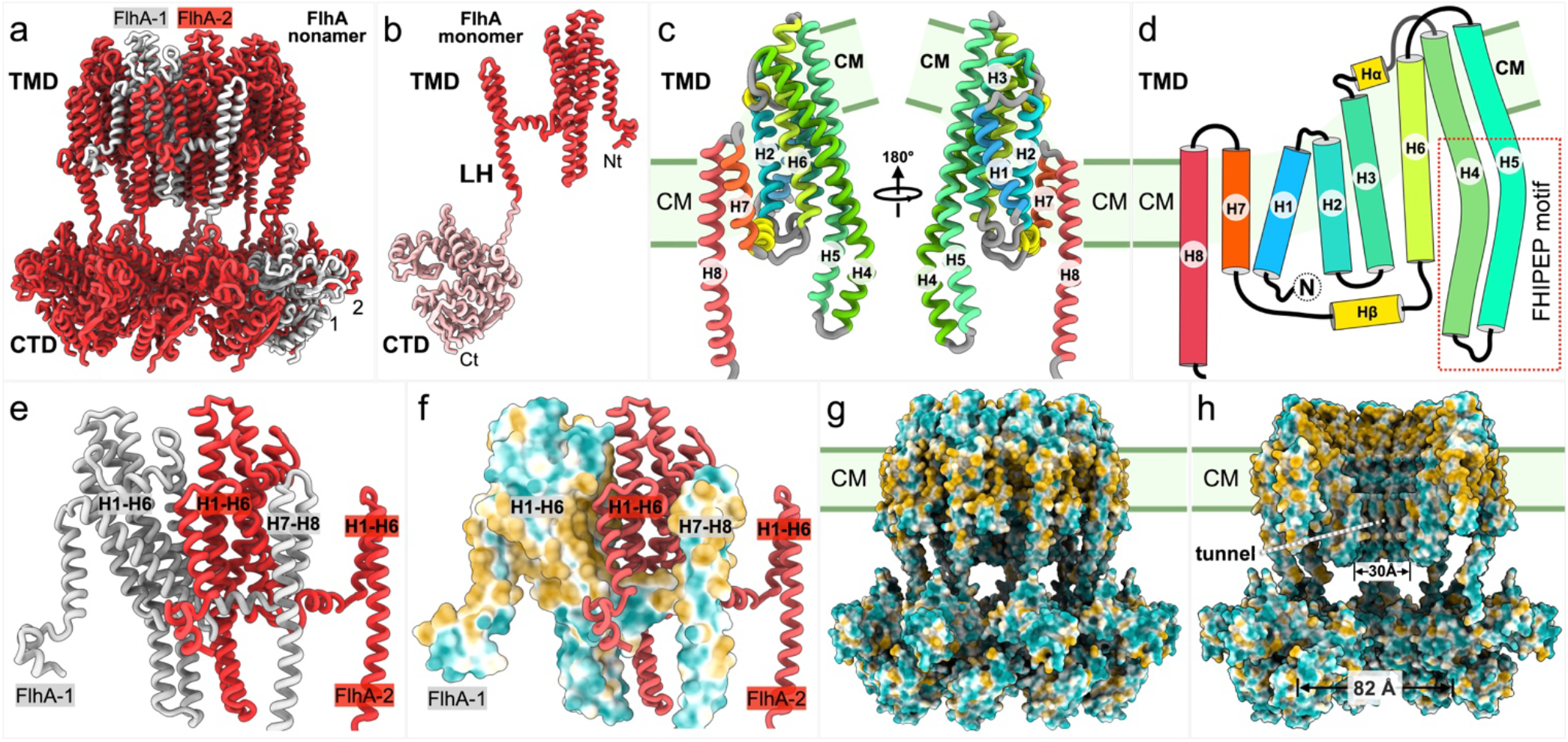
*In-situ* structure of the FlhA nonamer. (a) FlhA forms a nonameric assembly, with one subunit highlighted. (b) Domain organization of FlhA, including the FlhA_TMD_, FlhA_LH_, and FlhA_CTD_. (c,d) The FlhA_TMD_ adopts a stepped arrangement of helices (H1-H8) within the membrane. (e,f) Adjacent subunits exhibit domain swapping stabilized primarily by hydrophobic interactions. (g,h) Surface representations reveal a hydrophobic inner tunnel within the FlhA ring.

Nonamer assembly is driven by extensive inter-subunit hydrophobic interactions (Fig. 3e,f). Helices H7-H8 engage neighboring protomers in a domain-swapped arrangement, positioning FlhA_CTD_ beneath the FlhA_TMD_ of a non-adjacent subunit (Fig. 3a). Helix H8 extends into the linker helix, connecting FlhA_TMD_ to FlhA_CTD_. Consistent with species-dependent variability in this linker, the FlhA_CTD_ nonamer exhibits conformational flexibility relative to the FlhA_TMD_ nonamer, including subtle rotational and vertical displacements shown in 3D classification (Extended Data Fig. 5g).

A defining feature of FlhA_TMD_ is the pronounced bending of helices H4 and H5, which line the inner wall of the nonamer. Their upper segments are hydrophobic and tilt outward (∼27°), forming a barrel-like scaffold that geometrically complements the cone-shaped FliPQR–FlhB export gate. By contrast, the lower segments of H4 and H5 are enriched in polar and charged residues, corresponding to the conserved FHIPEP motif^38^ (Fig. 3). These residues define a central hydrophilic tunnel ∼30 Å in diameter, ideally suited for substrate translocation (Fig. 3).

### Asymmetric, lipid-mediated coupling between FlhA and the FliPQR–FlhB export gate

Our *in-situ* structure of the FliPQR–FlhB export gate from wild-type *B. burgdorferi* shows that, surprisingly, the M-gate occludes the central channel (Fig. 4a), similar to the closed conformation of the purified export gate from *Vibrio mimicus*^17,18^. This finding suggests that, in wild-type cells bearing fully assembled flagella, the export apparatus adopts a closed state, preventing type III secretion and flagellar assembly. FlhB contributes four transmembrane helices arranged as two hairpins that pack against the exterior of the FliPQR complex, while intervening loops wrap around the base of the gate (Fig. 4a-c; Extended Data Fig. 9a,b,c,e). In addition, an extended FlhB segment (Asp240-Met265), absent from previous models^17,18^, decorates the gate entrance and displaces the substrate entry site away from the central axis of the FlhA_TMD_ nonamer (Fig. 4e,f; Extended Data Fig. 9f-h). This offset entrance is surrounded by conserved basic residues in FlhA (Lys143, Arg147, Arg154, and Lys203), generating a strongly electropositive environment that likely promotes substrate recruitment and translocation (Fig. 4d,e). Consistent with this interpretation, mutations at equivalent positions in *S. enterica* severely impair substrate transport and bacterial motility^39,40^.

**Fig. 4:**
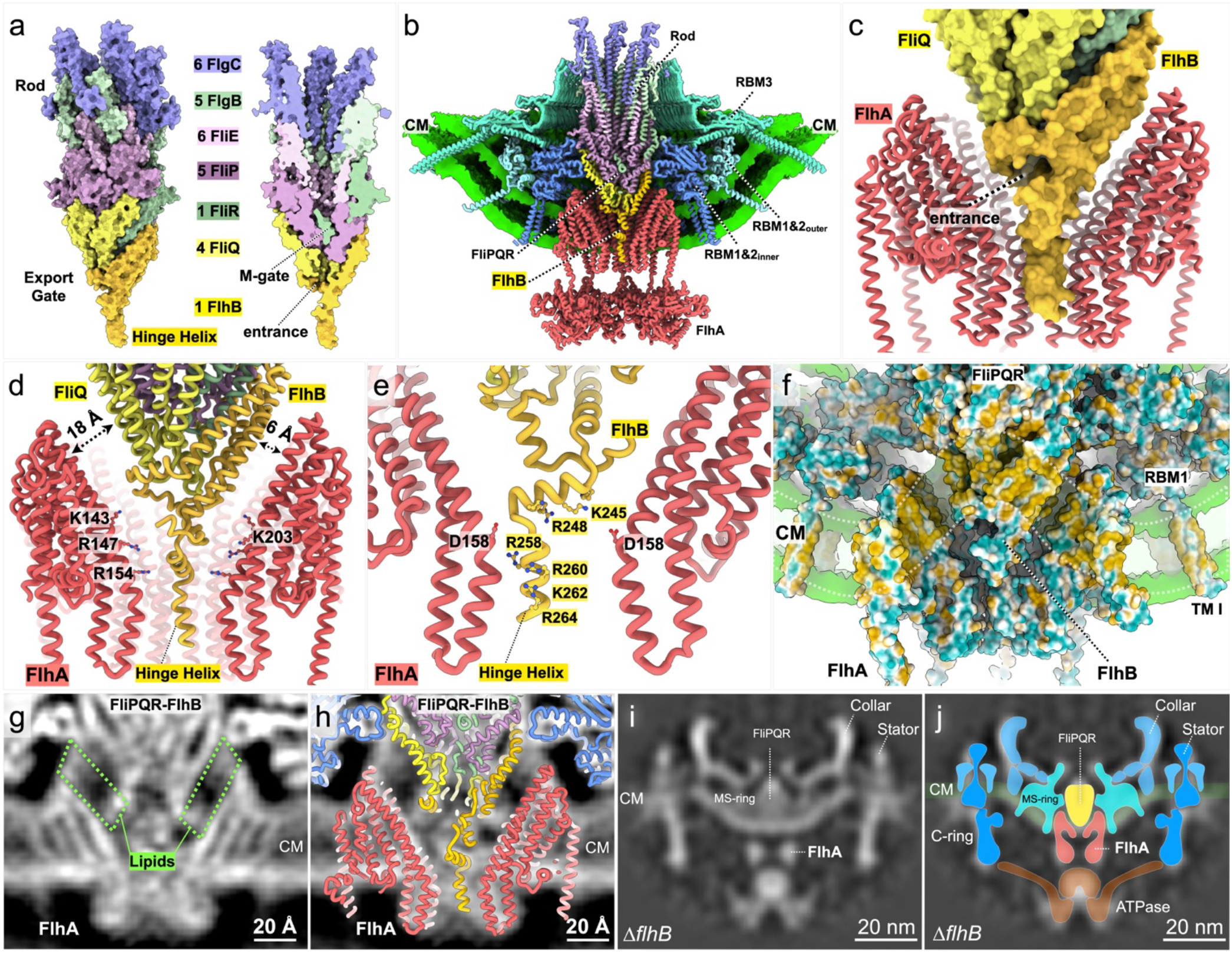
Lipid-mediated and asymmetric coupling between FlhA and the export gate. (a) Structure of the FliPQR–FlhB export gate in a closed conformation, with the M-gate occluding the channel. (b,c) Central sections reveal asymmetric positioning of the substrate entry site relative to FlhA. (d) FlhA and the export gate are separated by an asymmetric gap. (e) The FlhB hinge helix is enriched in basic residues and asymmetrically positioned along the inner tunnel wall. (f-h) Lipid-like densities occupy the gap between FlhA and the export gate, suggesting lipid-mediated coupling. (i,j) Cryo-ET analysis of a Δ*flhB* mutant shows that FlhA assembly remains in the absence of FlhB.

Notably, the FliPQR–FlhB export gate and FlhA do not make direct contact within the cytoplasmic membrane. Instead, a pronounced asymmetric gap separates the two assemblies, with a wider separation on the FliQ-facing side (∼18 Å) and a narrower distance on the FlhB-facing side (∼7 Å) (Fig. 4d). The gap is occupied by a thin, asymmetric bilayer-like density, forming a locally hydrophobic environment that conforms to both assemblies (Fig. 4f-h). Given that the central rod and export gate rotate in the wild-type flagellar motor, this lipid-mediated interface may provide a flexible yet sealed coupling between the export gate and FlhA, well suited to accommodate rotation.

Although the transmembrane interface is primarily lipid mediated, a direct interaction may occur within the FlhA hydrophilic tunnel. The hinge helix of FlhB (Thr240-Met265), enriched in arginine and lysine residues, is asymmetrically positioned along one side of the tunnel wall. This arrangement suggests that the FlhB hinge helix contributes to functional coupling between FlhA and the export gate during secretion. Consistent with this model, deletion of *flhB* does not disrupt FlhA nonamer assembly but abolishes rod, hook, and filament formation, as revealed by the *in-situ* motor structure of the Δ*flhB* mutant (Fig. 4i,j), thereby uncoupling assembly from export. Furthermore, the lipid mediated export apparatus creates a “funnel”-like conduit which narrows progressively from the cytoplasmic and transmembrane domains of the FlhA nonamer toward the entrance of the export gate (Fig. 4b), potentially facilitating effective substrate translocation.

### A hydrophilic pathway across FlhA may provide a route for proton translocation

Our *in-situ* structures reveal a continuous hydrophilic pathway lined by polar and charged residues between helices H2-H6 within FlhA and spanning the inner membrane from the periplasmic to cytoplasmic side (Fig. 5a,b). This feature appears conserved in AlphaFold3-predicted FlhA homologs (Extended Data Fig. 5), suggesting it represents a general property of T3SSs. Together with prior evidence that disruption of the ATPase complex does not abolish export in *B. burgdorferi*^41,42^, these results are consistent with a model in which FlhA mediates PMF-driven protein export^8,9^.

**Fig. 5:**
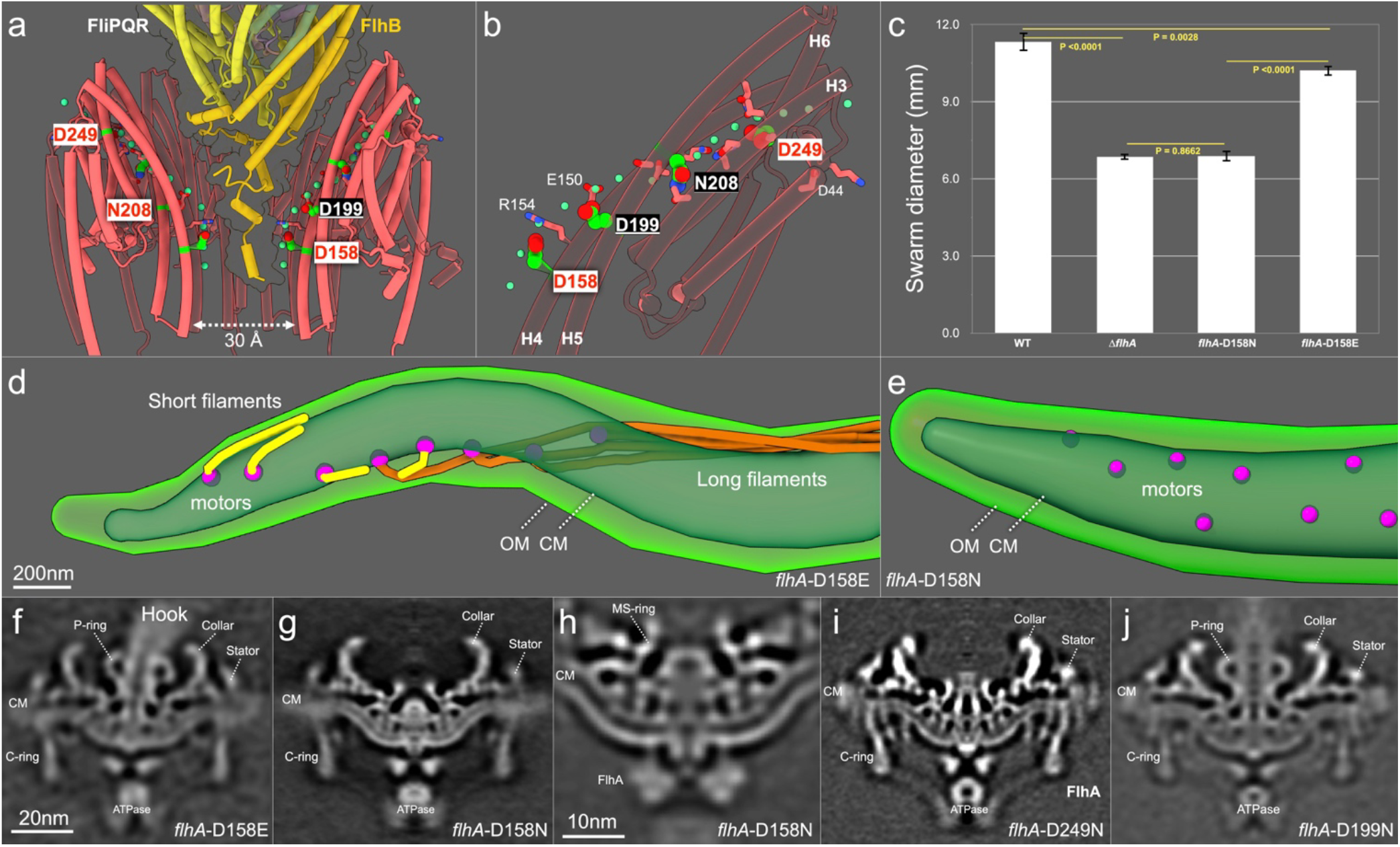
A hydrophilic pathway in FlhA underlies proton-coupled export. (a,b) Central sections reveal a continuous hydrophilic pathway within FlhA lined by conserved polar residues. (c) Mutational analysis of *flhA*-D158E, *flhA*-D158N, and Δ*flhA* demonstrates impaired motility. (d,e) Cryo-EM images reveal shorter flagellar filaments in the *flhA*-D158E mutant and no detectable flagellar filaments in the *flhA*-D158N mutant. (f-i) *In-situ* motor structures show that the *flhA*-D158E mutant assembles the rod and hook, whereas the *flhA*-D158N and *flhA*-D249N mutants lack both structures. (j) The in-situ motor structure from the *flhA*-D199N mutant shows the rod and the P-ring.

To better understand the roles of the continuous hydrophilic pathway within FlhA, we mutated conserved acidic residues (Asp158, Asp199, and Asp249). Asp158, located along the inner wall of the FlhA tunnel and oriented toward the FlhB hinge helix, is critical for function: *flhA*-D158E cells showed reduced motility and produced short flagella (Fig. 5c,d,f), whereas *flhA*-D158N cells were non-motile and lacked rod-hook-filament structures (Fig. 5c,e,g). Asp249, positioned near the periplasmic end (Fig. 5a,b), is also essential, as the *flhA*-D249N mutant abolished motility and rod-hook-filament assembly while retaining the overall architecture of the export apparatus (Fig. 5i). By contrast, Asp199, located above Asp158 in the tunnel, is dispensable, as *flhA*-D199N cells remained motile and showed no detectable structural defects in the flagellar motor (Fig. 5j). Notably, a conserved Asp208 is replaced by Asn208 in *B. burgdorferi* and *T. pallidum*, suggesting species-specific tuning of the pathway while preserving its polar character (Extended Data Fig. 8). Together, these data identify critical acidic residues along the hydrophilic pathway in FlhA that likely contribute to proton translocation and support PMF-driven protein export.

## DISCUSSION

Bacterial T3SSs assemble flagella or secrete virulence factors through a conserved transmembrane export apparatus, yet its organization and function in native membranes have remained unresolved at the molecular level^5-7^. Here, by integrating *in-situ* cryo-EM, cryo-ET, genetics, and modeling, we define the architecture of the membrane-bound export apparatus and surrounding MS-ring and reveal the molecular mechanisms underlying their assembly, energy transduction, and type III secretion.

T3SS-mediated flagellar assembly has been directly visualized in *B. burgdorferi*^35^ and other bacteria^11,12,43^, yet the precise sequence and interdependence of the export apparatus and MS-ring during the initial stages of flagellar motor formation remained unclear. Two models have been proposed: (i) an MS-ring-first pathway, in which FliF oligomerization in the cytoplasmic membrane precedes recruitment of the export apparatus^44^ and (ii) an export apparatus-first pathway, in which the transmembrane FliPQR–FlhAB complex nucleates subsequent MS-ring formation^45,46^. Our data show that FliF can assemble into the MS-ring in the absence of the export apparatus, albeit with very low efficiency. Strikingly, MS-ring assembly without the export gate or its core components (FliP, FliQ, and FliR) induces a U-shaped membrane deformation, distinct from the characteristic W-shaped membrane architecture observed in its presence, thereby preventing the transition between these two conformations. Importantly, our structural and genetic analyses demonstrate that formation of the unique export apparatus architecture strictly depends on a sequential, coordinated assembly process: the membrane-bound export gate nucleates the system, FlhA is subsequently recruited, and FliF oligomerization drives local membrane deformation while elevating the export apparatus to generate a funnel-shaped architecture (Fig. 6). This funnel-shaped conduit progressively narrows from the cytoplasmic ATPase complex to the FlhA cytoplasmic and transmembrane domains and finally to the export gate. This narrowing integrates substrate recruitment, chaperone coordination, and protein translocation within a preformed conduit across the membrane.

**Fig. 6:**
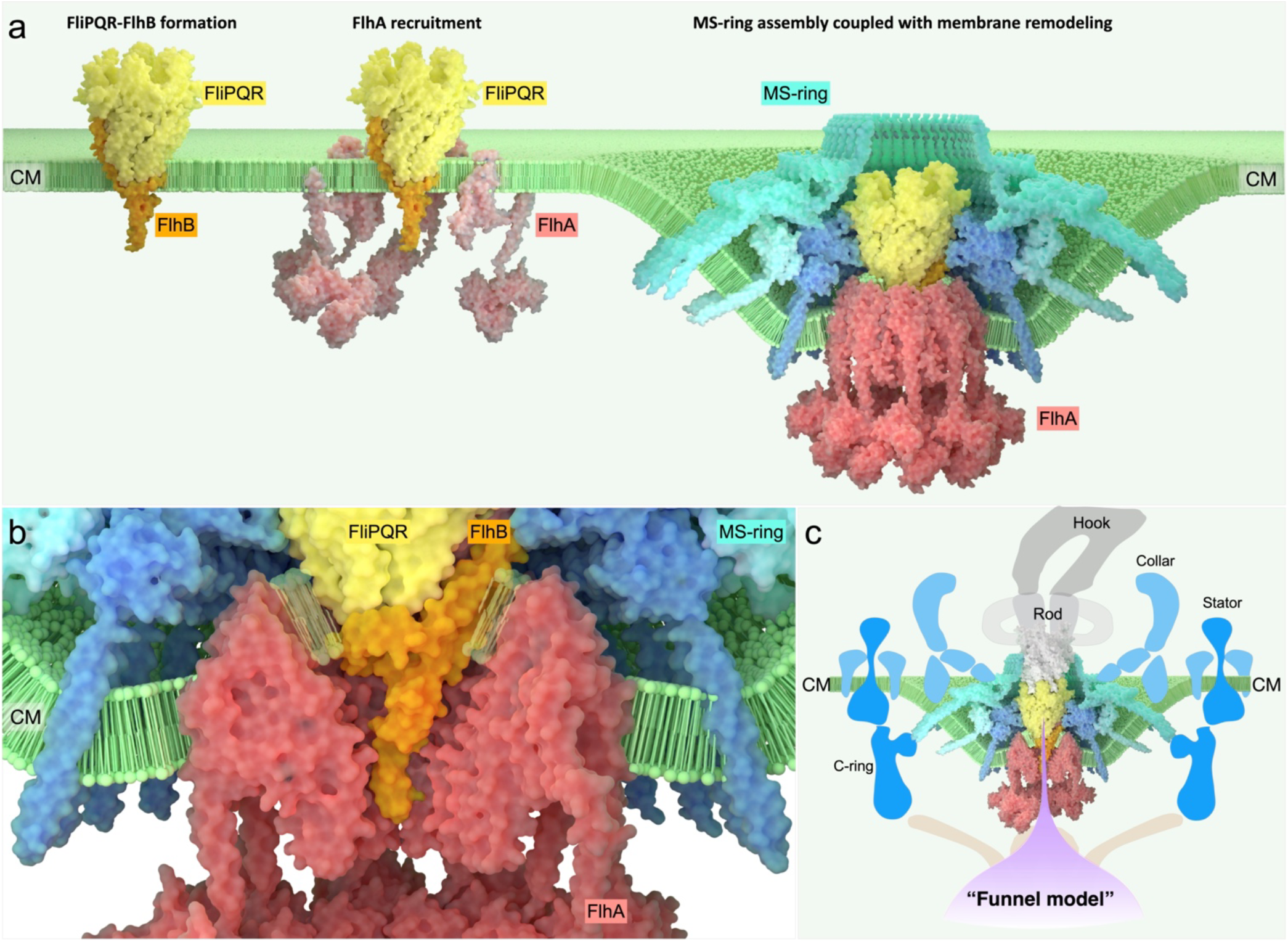
Model for assembly and energy coupling in the flagellar export apparatus. (a) Sequential, coordinated assembly model: FliPQR–FlhB assembles first, followed by FlhA recruitment and FliF incorporation, collectively driving progressive deformation of the membrane into a W-shaped architecture. (b) Zoom-in view of the export apparatus embedded in the curved membrane shows detailed protein-lipid-protein interactions. (c) Integrated model of the export apparatus as a funnel-shaped machine: the ATPase complex (∼60 nm) recruits and unfolds substrates, FlhA forms ∼5 nm and ∼2 nm chambers, and the FliPQR gate forms a narrow secretion channel.

We further identify a conserved hydrophilic pathway within FlhA, critically positioned within the funnel, that plays an essential role in PMF-mediated protein export. The positioning of key acidic residues near the FlhB hinge helix suggests a mechanism whereby proton flow through FlhA is coupled to opening of the export gate, linking PMF utilization to substrate translocation. Strikingly, this system lacks direct transmembrane contacts between the FlhA nonamer and export gate. Instead, these assemblies are separated by a thin lipid bilayer, indicating lipid-mediated coupling. This arrangement may preserve membrane integrity while allowing the flexibility required for flagellar rotation and dynamic conformational changes, highlighting the membrane as a key player in export apparatus assembly and type III secretion.

Together, these findings support a model in which the flagellar T3SS operates as a membrane-embedded, funnel-shaped nanomachine that integrates membrane deformation with energy transduction to achieve efficient protein export^12^. The strong conservation of this architecture suggests that these principles extend broadly across flagella and related injectisomes, providing a foundation for understanding type III secretion essential for bacterial motility and pathogenesis.

## MATERIALS AND METHODS

### Bacterial strains and growth conditions

High-passage *B. burgdorferi* strain B31A (WT) and its isogenic mutants (Extended Data Table 1) were grown in BSK-II liquid medium supplemented with 6% rabbit serum or on semi-solid agar plates at 35°C in the presence of 3% carbon dioxide as previously described^47,48^.

### Construction of Δ*fliP*, Δ*fliQ*, Δ*fliR*, Δ*flhA*, Δ*flhB* and four *flhA* point mutants

Single mutants were generated using a gene inactivation methodology designed to create deletion mutants while avoiding any polar effects on downstream gene expression^48^. For instance, the *fliP* gene (gene locus BB0275; a 765 bp gene) was inactivated by replacing *fliP* with the *aadA* streptomycin/spectinomycin resistance gene coding sequence through overlapping PCR. The process involved three steps of DNA amplification using PCR. In the first step, each DNA region was amplified individually using PCR primer pairs P1-P2 (5′-upstream *fliP*, 3′-upstream *fliP*), P3-P4 (aadA coding sequence), and P5-P6 (5′-downstream *fliP*, 3′-downstream *fliP*). Primers P2, P3, P4, and P5 contained overlapping regions. In the second step, a PCR product was generated using primers P1 and P4 and purified DNA products for upstream *fliP* and aadA as templates. In the third step, the final PCR product was obtained using primers P1 and P6, and the purified DNA products of upstream *fliP*-aadA and downstream *fliP* served as templates to amplify the upstream *fliP*-aadA-downstream *fliP* DNA construct.

The resulting PCR product was 2,462 bp in length and was subsequently purified from a gel and cloned into the pGEM-T Easy vector (Promega Inc.). The integrity of the *fliP* inactivation plasmid was confirmed through PCR and restriction mapping. Electroporation of PCR-amplified DNA into B31A competent cells followed by plating on BSK-II medium containing 100 μg/ml streptomycin. Construction of **Δ***fliQ*, **Δ***fliR*, **Δ***flhA*, and **Δ***flhB* was performed in the same strategy mentioned above for **Δ***fliP*.

Point mutations of *flhA-*D158E, *flhA-*D158N, *flhA*-D199N, and *flhA*-D249N were generated using a promoterless kanamycin cassette and a site-directed point mutagenesis kit, as previously described^49,50^. The mutants were confirmed through PCR and DNA sequencing.

### Construction of a quintuple mutant Δ*fliP-flhA*

Deletion of the *fliP, fliQ, fliR, flhB*, and *flhA* genes was accomplished utilizing the Cre-lox recombination system^51^. Initially, LoxP sites were introduced into the *fliP* and *flhA* genes on the chromosome in the same orientation, leading to the deletion of the sequence containing these genes, as well as *fliQ, fliR*, and *flhB*. The introduction of LoxP sites was achieved by PCR amplification of the *fliP* and *flhA* genes, each containing a single HindIII site. The PCR products, yielding a 744-bp fragment for *fliP* and a 1306-bp fragment for *flhA*, were gel purified and cloned into the pGEM-T Easy vector (Promega Inc.). Subsequently, a HindIII restriction site was engineered to flank the loxP site along with streptomycin resistance (pABA07) using PCR primer pairs P35-36, and the resulting fragment was cloned into the pGEM-T Easy vector. The loxP site with the streptomycin resistance cassette in pGEM-T Easy and the loxP site with a kanamycin resistance cassette (pABA14) were digested with HindIII and cloned into plasmids containing the *fliP* and *flhA* genes in pGEM-T Easy, respectively. These plasmids were also digested with HindIII, resulting in the creation of the loxP insertion mutant vectors. The integrity of the loxP insertion mutant vectors and the orientation of the loxP sites were confirmed through PCR and restriction mapping.

### Determination of polar effect on downstream gene expression

Even though our novel gene inactivation system does not impose any polar effects as we have confirmed and verified previously^35,52^, we still determined the effect of a deletion mutant on the expression of the downstream genes using qRT-PCR as described previously^53^. Total RNA was extracted from exponentially-grown *B. burgdorferi* (10 ml) cells by using Direct-zol™ RNA MiniPrep Kit (Zymo Research). The RNA preparation was digested with Turbo DNase I (Ambion) overnight to ensure that the samples were free of contaminating genomic DNA. The concentration and purity of each RNA sample were measured via spectrophotometry (ND-1000 spectrophotometer; NanoDrop Technologies) and were also assessed by gel electrophoresis. Samples were checked for contamination of genomic DNA by PCR, using *B. burgdorferi* enolase primers. First-strand cDNA was prepared using the AffinityScript cDNA Synthesis Kit (Agilent) according to the manufacturer’s instructions. The resulting cDNA was amplified using a CFX96 Real-Time System (Bio-Rad), with a final reaction volume of 25 μl that contained 10 ng of cDNA, Power SYBR® Green PCR Master Mix (Life Technologies), and *B. burgdorferi* gene-specific primers. *B. burgdorferi* enolase was used as an internal control. Real-time PCRs were carried out in triplicate, and the data indicate that polar effects on the downstream genes did not occur (data not shown).

Sodium dodecyl sulfate-polyacrylamide gel electrophoresis (SDS-PAGE) followed by Western blotting was performed using a Thermo Scientific SuperSignal kit, as described^54^. The protein concentration in the cell lysates was measured using a Bio-Rad protein assay kit. Approximately 20 μg of lysates were loaded into each lane of the SDS-PAGE gel, and the samples were subjected to either coomassie blue staining or immunoblotting with specific antibodies targeting *B. burgdorferi* FliF and DnaK^**54**^.

### Dark-field microscopy and swarm plate assays

Exponentially growing *B. burgdorferi* cells were observed using a Zeiss Axio Imager M1 dark-field microscope to visualize the growth, morphology, and motility. Swarm plate assay was performed to determine spirochetes’ motility using our established protocol^55^.

### Frozen-hydrated cryo-EM sample preparation

The preparation of frozen-hydrated specimens followed established protocols^56^. Briefly, *B. burgdorferi* cultures were subjected to centrifugation at 5,000 ×g for 5 minutes. The resulting pellet was resuspended in 1.0 ml of PBS, followed by another centrifugation step and suspension in ∼50 μl of PBS. For cryo-ET and cryo-EM, the cell suspension was mixed with or without 10 nm colloidal gold fiducial markers. A 5 μl aliquot of the cell suspension (OD_600_≈0.2) was applied to freshly glow-discharged holey carbon grids (Quantifoil) or lacey carbon grids (Ted Pella). Grids were blotted for 6–8 s using filter paper and rapidly plunge-frozen in liquid ethane using a GP2 plunger (Leica).

### Cryo-ET data collection and analyses

Frozen-hydrated specimens were imaged at -170°C using a 300 kV Krios electron microscope (Thermo Fisher Scientific) equipped with a field emission gun, Volta phase plate, and a post-GIF direct electron detector (Gatan) with a magnification of ×42,000, resulting in a pixel size of 2.148 Å. Tilt series were collected from −48° to 48° with 3° increments using SerialEM^57^. The total dose of the tilt series was ∼70 e^-^Å^2^ at a defocus of -4.8 µm. Frozen-hydrated specimens of four mutants were imaged at -170° C using a Polara G2 electron microscope (FEI) equipped with a field emission gun and a 4K×4K CCD camera (TVIPS). The microscope was operated at 300 kV with a magnification of ×31,000, resulting in an effective pixel size of 5.7 Å after 2×2 binning. Low-dose, single-axis tilt series were collected from each cell at -6 to -8 µm defocus with a cumulative dose of ∼100 e-/Å2 distributed over 87 images and covering an angular range of -64° to +64°, with an angular increment of 1.5°. The number of tomograms utilized for each strain is detailed in Extended Data Table 2. In total, 3,302 tomograms were collected from 11 different *B. burgdorferi* strains.

Recorded images were initially motion-corrected using MotionCorr2^58^ and subsequently stacked by IMOD^59^. Tilt series alignment was performed using fiducial markers or fiducial-free alignment by IMOD^59^. Gctf^60^ was employed to determine the defocus of each tilt image in the aligned stacks, and the “ctfphaseflip” function in IMOD^59^ was used for contrast transfer function (CTF) correction of the tilt images. Tomograms were generated using Tomo3D^61^ by either simultaneous iterative reconstruction technique (SIRT) or weighted back-projection (WBP). SIRT reconstructions were utilized for particle picking of flagellar motors due to their higher contrast, while tomograms reconstructed by WBP retained high-resolution details for *in-situ* structure determination of the flagellar motor through subtomogram averaging.

### Subtomogram averaging and corresponding analysis

15,962 flagellar motors were manually picked from 3,403 tomographic reconstructions of 11 different *B. burgdorferi* strains (Extended Data Table 2). The protomo package was used to perform 3D alignment and classification as described previously^62,63^. Initially, motors were aligned within 6× binned tomograms based on the estimated initial orientation, utilizing the center coordinates of the flagellar C ring and collar to provide two of the three Euler angles. Subsequently, subtomograms were extracted from unbinned tomograms using the refined positions and 4× binned for alignment. This process was reiterated with 2× binned subtomograms to enhance structure detail.

Focused refinement was then employed to enhance the high resolution of the export apparatus and MS-ring structures. Multi-reference alignment and 3D classification were utilized^62,64^. Initially, the region around the MS-ring and export apparatus was extracted from each motor. Subsequently, 3D alignment refinement and 3D classification were applied to eliminate particles exhibiting poor contrast or significant distortions, resulting in refined structures. Fourier shell correlation was calculated by comparing two randomly divided halves of the aligned images, enabling estimation of the final maps’ resolution.

### *In-situ* single-particle cryo-EM data collection and preprocessing

Data were collected on a Titan Krios electron microscope (Thermo Fisher Scientific) operated at 300 kV and equipped with a K3 direct electron detector and a GIF energy filter (Gatan). Movies were recorded in super-resolution mode at a nominal magnification of ×81,000, corresponding to a calibrated pixel size of 0.534 Å. Automated data acquisition was performed in SerialEM using a customized multishot script to target bacterial cells directly on cryo-EM grids. Each movie was recorded with a total electron dose of 70 e^-^/Å^2^ fractionated over 40 frames, with a defocus range of −1.2 to −2.2 μm. Full data collection parameters are summarized in Extended Data Table 3. Motion correction and contrast transfer function (CTF) estimation were performed in CryoSPARC^65^.

### Flagellar particle selection and structure determination

All image processing and structural analyses were performed in CryoSPARC^65^, with the overall workflow summarized in Extended Data Fig. 3. To establish an automated particle-picking model, ∼1,000 flagellar motors were manually picked and used to train a neural-network model in Topaz^66^. The model was iteratively refined and subsequently applied for automated particle picking across the full dataset. In total, 26,472 micrographs were collected, yielding ∼87,807 particles after Topaz-based picking. Following 2D classification, ∼56,720 particles displaying clear motor features, including both top and side views, were retained for further analysis. Particles were extracted with a box size of 448 pixels at pixel size of 1.14 Å.

An initial 3D model was generated using ab-initio reconstruction with C16 symmetry. Subsequent heterogeneous refinement with C16 symmetry was used to further sort particle populations. Focused refinement on the whole basal-body region improved particle alignment of intact motors, resulting in a reconstruction from 43,970 particles at an overall resolution of ∼8 Å.

For detailed analysis of the export apparatus and MS-ring, signals corresponding to the stator and peripheral cage/collar regions were subtracted, followed by C16 symmetry expansion. The resulting particles were subjected to focused 3D classification on the MS-ring without alignment (C1 symmetry). This analysis revealed that the MS-ring is composed of 41 subunits, with the RBM3 domain forming the S ring with C41 symmetry. The RBM1 and RBM2 domains form two concentric assemblies: an inner ring with C23 symmetry and a more flexible outer ring with C18 symmetry. After removal of duplicated particles, 40,949 particles were retained for downstream refinement. Focused refinement of each structural module, combined with local CTF refinement, further improved local resolution.

To address symmetry mismatch between different components of the motor, particles were symmetry-expanded according to the refined MS-ring RBM3, followed by focused classification and local refinement with corresponding masks. The central rod was refined under C1 symmetry, whereas FlhA was refined under C9 symmetry. Focused refinement, combined with local CTF refinement further improved the rod and FlhA densities.

For the, although the FlhA_TMD_ was resolved at high resolution, the FlhA_CTD_ appeared diffuse, indicating conformational flexibility. To further characterize this variability, focused 3D classification and 3D variability analysis^67^ were performed with masks covering the FlhA_CTD_, enabling assessment of conformational heterogeneity.

Local resolution of each map was estimated in CryoSPARC^65^ using the gold-standard FSC 0.143 criterion, as summarized in Extended Data Fig. 3. Final EM maps were post-processed using EMReady^68^ to facilitate model building and visualization.

### Model building and refinement

Initial models were generated from AlphaFold3^69^-predicted monomer structures and rigid-body fitted into the EM density maps using UCSF ChimeraX^70^. Flexible fitting was subsequently performed using Namdinator^71^, followed by manual model adjustment in Coot^72^. Real-space refinement was carried out in PHENIX^73^ with geometry restraints. Model quality was assessed using MolProbity^74^.

## Data availability

All cryo-EM maps and atomic coordinates for the export apparatus and the MS-ring have been deposited in the Electron Microscopy Data Bank (EMDB) and the Protein Data Bank (PDB), including both overall and locally refined maps. The MS-ring RBM3 domain (C41 symmetry), EMD-76344/PDB: 12DR; the MS-ring RBM1&2 inner ring (C23 symmetry), EMD-76345/PDB: 12DS; the MS-ring RBM1&2 outer ring (C18 symmetry), EMD-76346/PDB: 12DT; the export gate–rod–RBM1&2inner complex (C1 symmetry), EMD-76347/PDB: 12DU; the FlhA transmembrane domain (C9 symmetry), EMD-76348/PDB: 12DV; and the overall structure of the export apparatus and MS-ring, EMD-76349/PDB: 12DW. The cryo-ET/STA map of the wild-type export apparatus, EMD-76383/PDB: 12EQ; the Δ*fliQ* export apparatus map, EMD-48081; and the Δ*flhA* export apparatus map, EMD-48082.

## ACKNOWLEDGMENTS

We thank Jennifer Aronson for critical reading of the manuscript and Kangkang Song for assisting in cryo-ET/EM data collection. We are grateful for the support from Drs. Jiagang Tu, Xiaowei Zhao, and Brittany Carroll. We are grateful for the antibodies provided by Dr. Chunhao (Chris) Li at Virginia Commonwealth University. The project was supported by grants R01AI087946, R01AI132818, R01AI078958 from the National Institute of Allergy and Infectious Diseases (NIAID) and the National Institutes of Health (NIH). Cryo-ET data were collected at the Yale CryoEM Resource, which is funded in part by NIH grant 1S10OD023603-01A1.

