## Supplemental Data for "*In-situ* structure of the flagellar export apparatus in *Borrelia burgdorferi*"

**Supplemental Information**


| | |
|---|---|
| Extended Data Fig. 1 | pg2 |
| Extended Data Fig. 2 | pg3,4 |
| Extended Data Fig. 3 | pg5,6 |
| Extended Data Fig. 4 | pg7,8 |
| Extended Data Fig. 5 | pg9 |
| Extended Data Fig. 6 | pg10 |
| Extended Data Fig. 7 | pg10 |
| Extended Data Fig. 8 | pg11,12 |
| Extended Data Fig. 9 | pg13 |
| Extended Data Table 1 | pg14 |
| Extended Data Table 2 | pg14 |
| Extended Data Table 3 | pg15,16 |

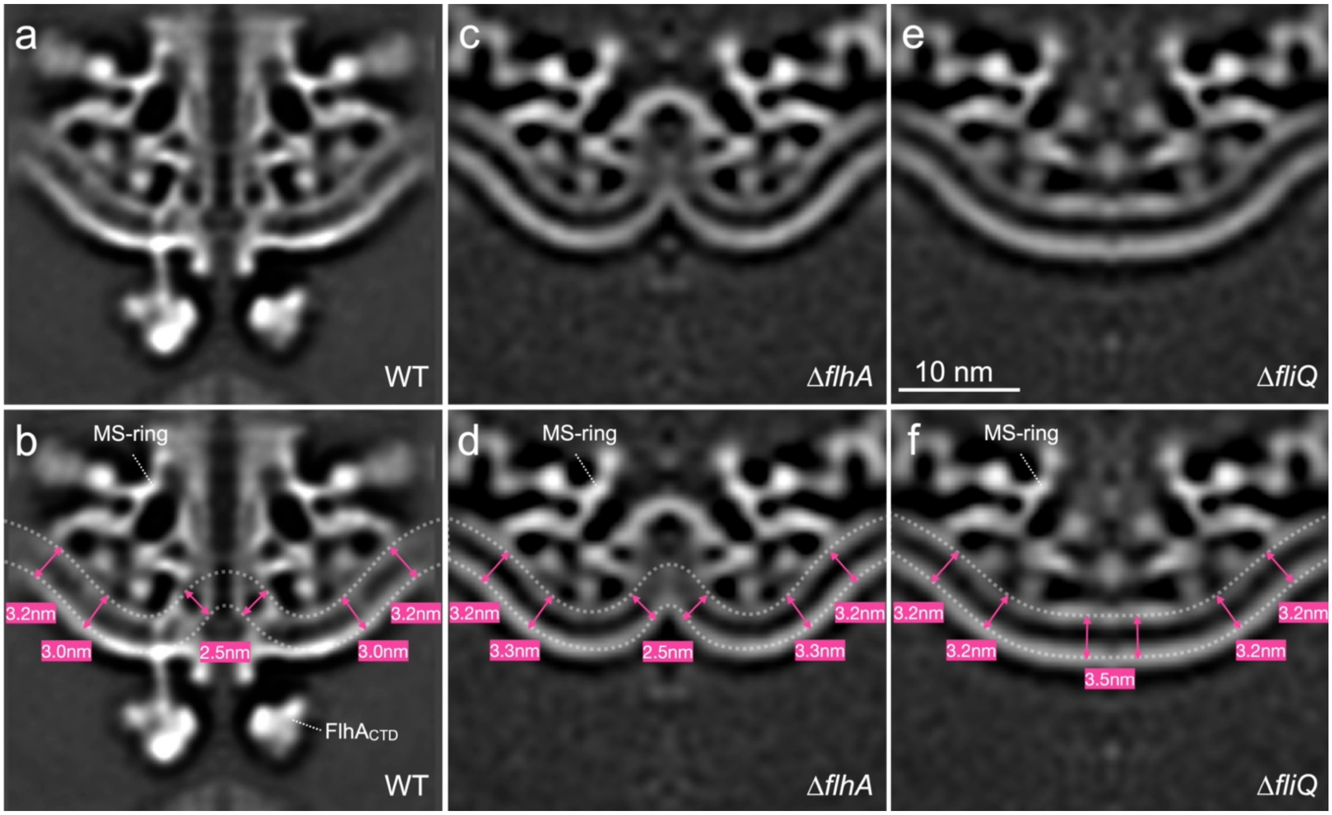


**Extended Data Fig. 1 | Membrane leaflet spacing beneath flagellar basal bodies in *B. burgdorferi*.** Central section (a) and spacing (b) between the two cytoplasmic membrane leaflets in the wild-type basal body, revealing variable spacing and pronounced membrane thinning near the export apparatus. (c, d) Corresponding views for the Δ*flhA* basal body. (e, f) Corresponding views for the Δ*fliQ* basal body.

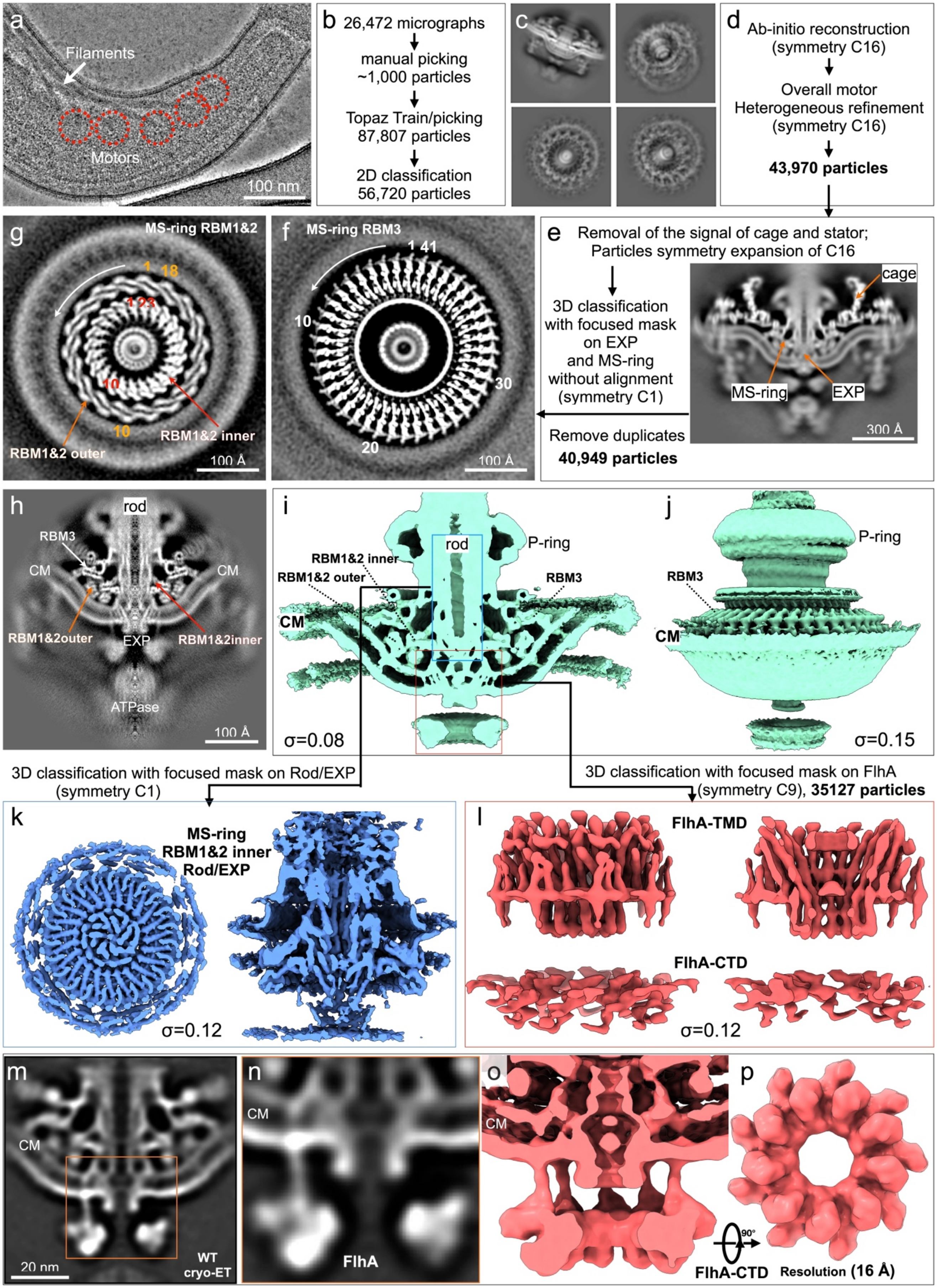
a
Filaments
Motors
100 nm
b 26,472 micrographs
manual picking
~1,000 particles
Topaz Train/picking
87,807 particles
2D classification
56,720 particles
c
d Ab-initio reconstruction
(symmetry C16)
Overall motor
Heterogeneous refinement
(symmetry C16)
43,970 particles
e Removal of the signal of cage and stator;
Particles symmetry expansion of C16
3D classification
with focused mask
on EXP
and MS-ring
without alignment
(symmetry C1)
cage
MS-ring
EXP
300 Å
Remove duplicates
40,949 particles
f MS-ring RBM3
1 41
10
30
20
100 Å
g MS-ring RBM1&2
1 18
1 2 3
10
10
RBM1&2 outer
RBM1&2 inner
100 Å
h rod
RBM3
CM
CM
RBM1&2outer
EXP
RBM1&2inner
ATPase
100 Å
i rod
P-ring
RBM1&2 inner
RBM1&2 outer
RBM3
CM
σ=0.08
j P-ring
RBM3
CM
σ=0.15
3D classification with focused mask on Rod/EXP
(symmetry C1)
3D classification with focused mask on FlhA
(symmetry C9), 35127 particles
k MS-ring
RBM1&2 inner
Rod/EXP
σ=0.12
l FlhA-TMD
FlhA-CTD
σ=0.12
m CM
20 nm
WT
cryo-ET
n CM
FlhA
o CM
90°
FlhA-CTD
p Resolution (16 Å)

**Extended Data Fig. 2 | Overall image processing and structure determination by in situ single-particle cryo-EM.** (a) Representative micrograph of a *B. burgdorferi* cell tip, with flagellar motors indicated. (b) Particle-picking workflow. From 26,472 micrographs, ~1,000 manually picked particles were used to train Topaz. Iterative picking and classification yielded 56,720 particles. (c) Representative 2D class averages. (d) 3D reconstruction workflow. Ab initio reconstruction (CryoSPARC, C16 symmetry), followed by heterogeneous refinement, yielded 43,970 particles for the overall structure (e, right). In parallel, removal of collar and stator densities, symmetry expansion (C16), and focused classification (C1, no further alignment) enabled analysis of the export apparatus (EXP) and MS ring (e, left). (f–h) Selected classes revealing MS-ring symmetries: C41 (RBM3), C23 (RBM1–RBM2 inner ring), and C18 (outer ring). (i, j) Surface-rendered MS-ring maps. (k) Focused classification on the central rod (no alignment), yielding classes with clear secondary-structure features. (l) Focused classification on FlhA (no further alignment), followed by local refinement and local CTF refinement. (m–p) In situ map of the FlhA cytoplasmic domain (CTD) (16 Å resolution) resolved by cryo-ET.

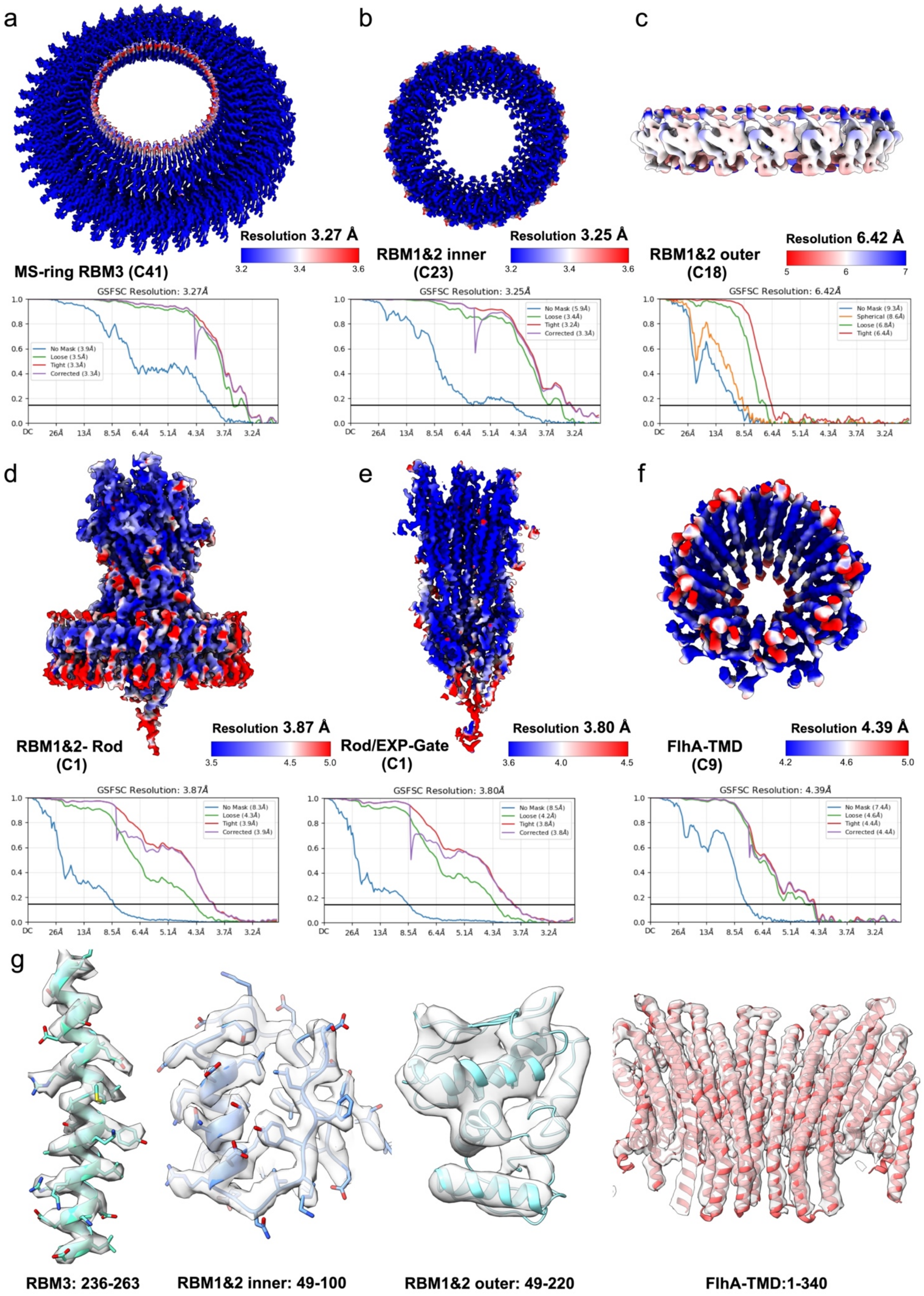
a
b
c
Resolution 3.27 Å
3.2
3.4
3.6
MS-ring RBM3 (C41)
RBM1&2 inner
(C23)
Resolution 3.25 Å
3.2
3.4
3.6
RBM1&2 outer
(C18)
Resolution 6.42 Å
5
6
7
GSFSC Resolution: 3.27Å
No Mask (3.9Å)
Loose (3.5Å)
Tight (3.3Å)
Corrected (3.3Å)
GSFSC Resolution: 3.25Å
No Mask (5.9Å)
Loose (3.4Å)
Tight (3.2Å)
Corrected (3.3Å)
GSFSC Resolution: 6.42Å
No Mask (9.3Å)
Spherical (8.6Å)
Loose (6.8Å)
Tight (6.4Å)
DC
26Å
13Å
8.5Å
6.4Å
5.1Å
4.3Å
3.7Å
3.2Å
d
e
f
RBM1&2- Rod
(C1)
Resolution 3.87 Å
3.5
4.5
5.0
Rod/EXP-Gate
(C1)
Resolution 3.80 Å
3.6
4.0
4.5
FlhA-TMD
(C9)
Resolution 4.39 Å
4.2
4.6
5.0
GSFSC Resolution: 3.87Å
No Mask (8.3Å)
Loose (4.3Å)
Tight (3.9Å)
Corrected (3.9Å)
GSFSC Resolution: 3.80Å
No Mask (8.5Å)
Loose (4.2Å)
Tight (3.8Å)
Corrected (3.8Å)
GSFSC Resolution: 4.39Å
No Mask (7.4Å)
Loose (4.6Å)
Tight (4.4Å)
Corrected (4.4Å)
g
RBM3: 236-263
RBM1&2 inner: 49-100
RBM1&2 outer: 49-220
FlhA-TMD:1-340

**Extended Data Fig. 3 | Resolution estimation and model fitting of individual subcomplex maps.** (a–c) MS-ring subcomplexes: RBM3 (C41 symmetry), RBM1–RBM2 inner ring (C23 symmetry), and RBM1–RBM2 outer ring (C18 symmetry). (d, e) Export gate–rod assemblies (C1 symmetry). (f) FlhA transmembrane domain (TMD; C9 symmetry). (g) Representative atomic models fitted into the corresponding density maps.

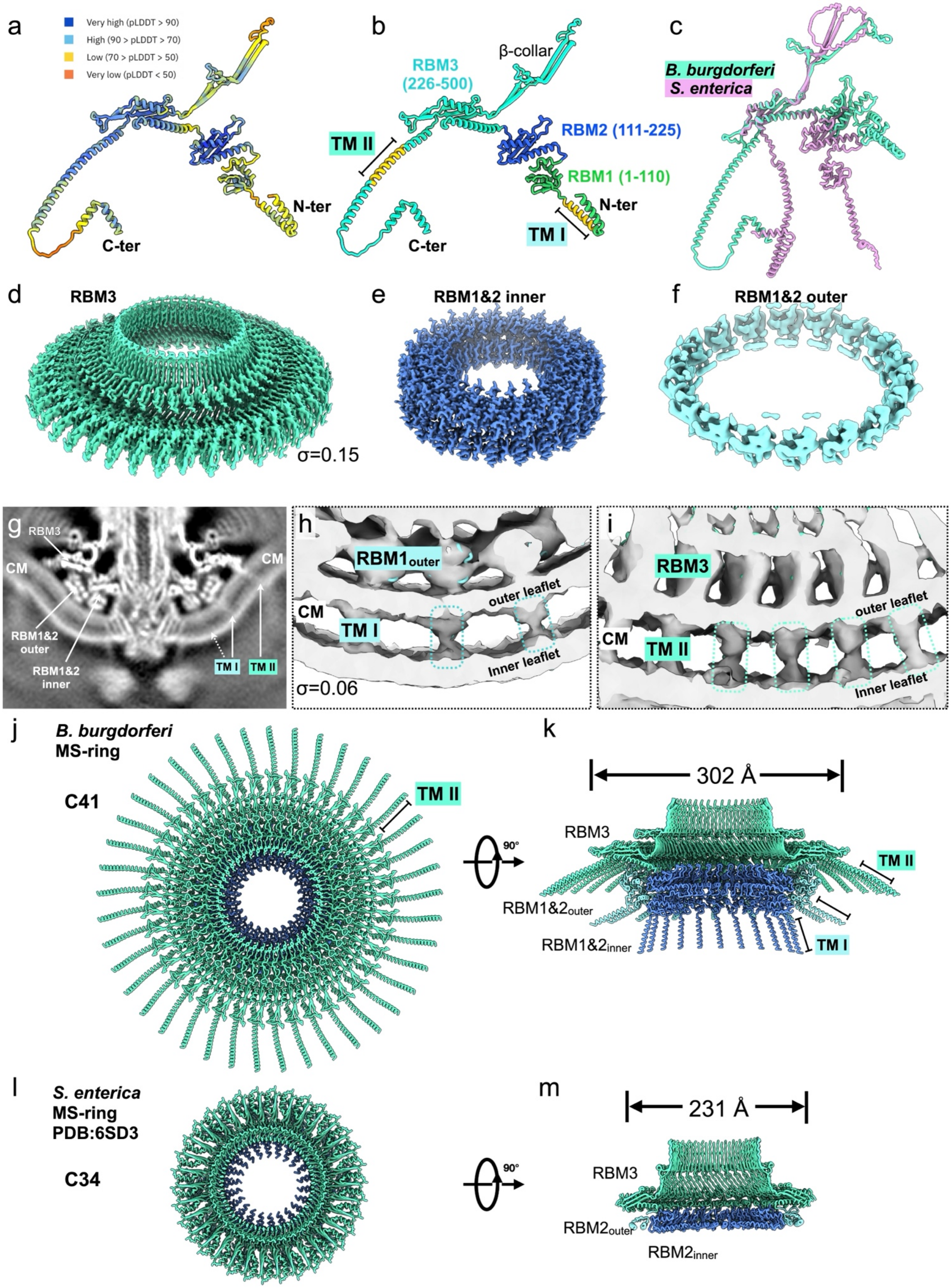
a
Very high (pLDDT > 90)
High (90 > pLDDT > 70)
Low (70 > pLDDT > 50)
Very low (pLDDT < 50)
N-ter
C-ter
b
RBM3 (226-500)
β-collar
RBM2 (111-225)
RBM1 (1-110)
TM II
TM I
N-ter
C-ter
c
B. burgdorferi
S. enterica
d
RBM3
σ=0.15
e
RBM1&2 inner
f
RBM1&2 outer
g
RBM3
CM
CM
RBM1&2 outer
RBM1&2 inner
TM I
TM II
h
RBM1outer
CM
TM I
outer leaflet
Inner leaflet
σ=0.06
i
RBM3
CM
TM II
outer leaflet
Inner leaflet
j
B. burgdorferi
MS-ring
C41
TM II
k
302 Å
90°
RBM3
TM II
RBM1&2outer
RBM1&2inner
TM I
l
S. enterica
MS-ring
PDB:6SD3
C34
m
231 Å
90°
RBM3
RBM2outer
RBM2inner

**Extended Data Fig. 4 | Structural comparison of the MS ring.** (a) AlphaFold3-predicted structure of the *B. burgdorferi* FliF monomer. (b) Domain organization of FliF. (c) Structural comparison of FliF from *B. burgdorferi* and *S. enterica*, showing strong conservation at the monomer level. (d) Cryo-EM structure of the RBM3 domain. (e) Cryo-EM structure of the inner rings of the RBM1&2 domains. (f) Cryo-EM structure of the outer ring of the RMB1&2 domains. (g) Cross-section of the cryo-EM structure. (h, i) Surface renderings show the transmembrane densities corresponding to TMI and TM2, respectively. (j, k) Top and central cross-sectional views of the *B. burgdorferi* MS ring. (l, m) Corresponding views of the *S. enterica* MS ring (PDB: 6SD3). Differences in stoichiometry (C41 versus C34 symmetry) result in distinct ring dimensions.

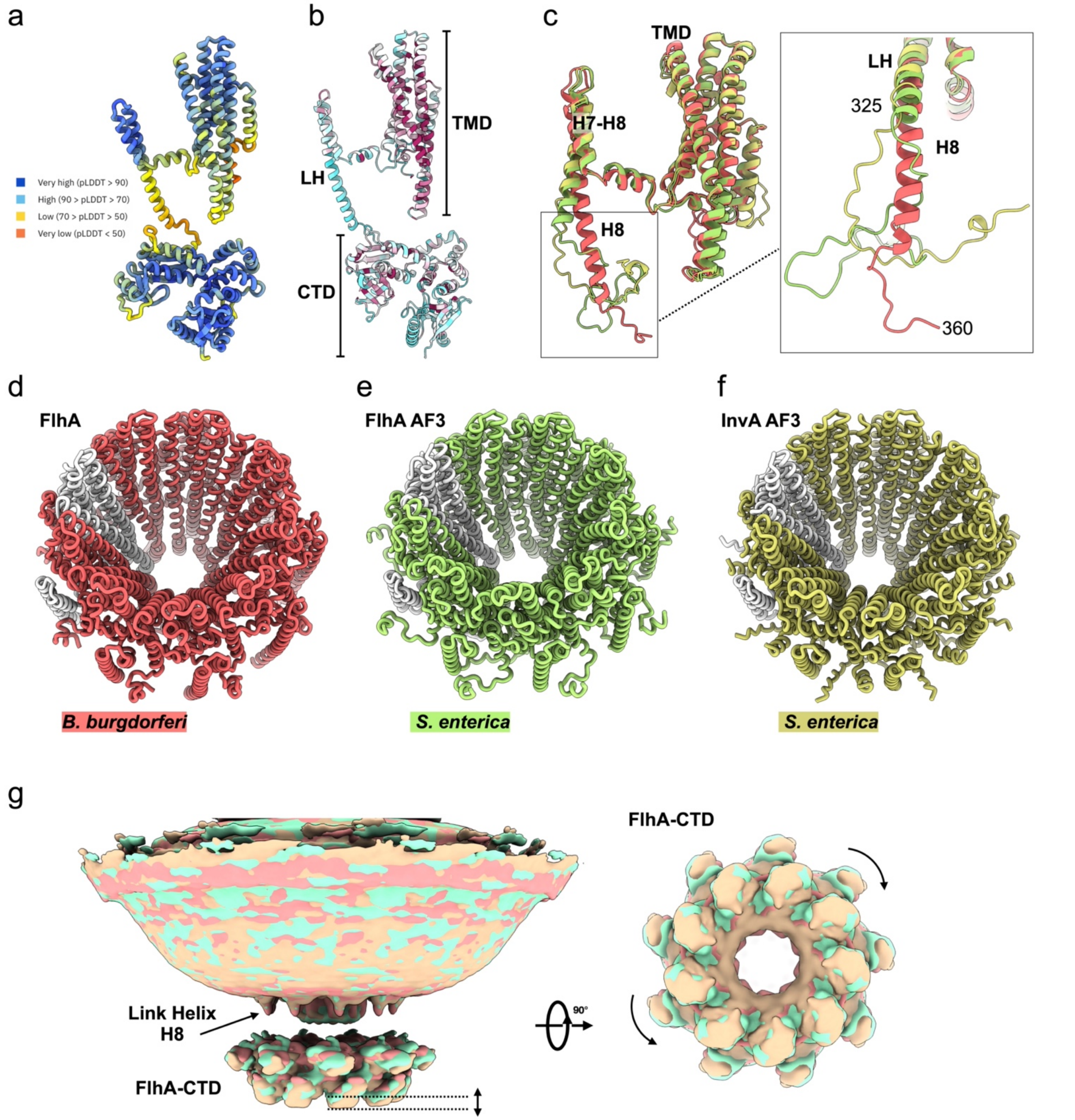


**Extended Data Fig. 5 | Comparison of FlhA structures.** (a) AlphaFold-predicted structure of the *B. burgdorferi* FlhA monomer. (b) Domain organization of FlhA, colored by sequence conservation. (c) Structural comparison of FlhA from *B. burgdorferi* and *S. enterica*, and InvA from the *S. enterica* injectisome. The transmembrane domain (TMD) is highly conserved, whereas the linker helix varies, adopting either helical or loop conformations. (d–f) Comparison of nonameric assemblies of *B. burgdorferi* FlhA and AlphaFold-predicted *S. enterica* FlhA and InvA. (g) 3D classification reveals conformational flexibility of the FlhA cytoplasmic domain (CTD), including vertical displacement and rotational variation, likely mediated by the linker helix.

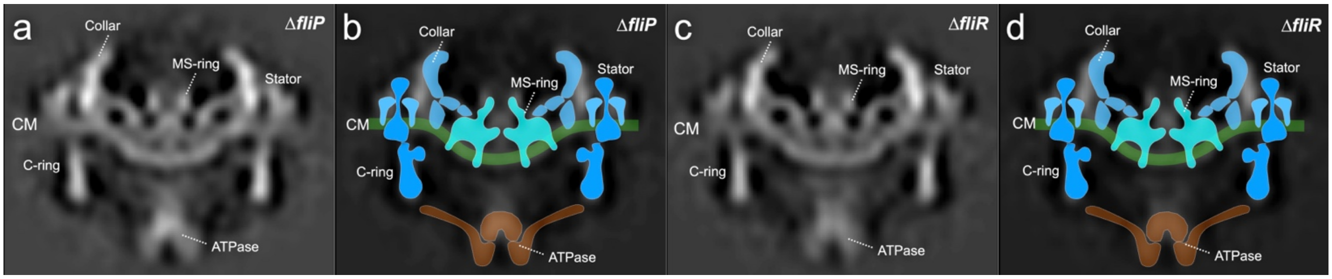


**Extended Data Fig. 6 | Characterization of the export apparatus in Δ*fliP* and Δ*fliR* mutants.** (a, b) Central section (**a**) and corresponding model (**b**) of the basal body structure in the Δ*fliP* mutant, showing absence of FlhA density. (c, d) Central section (**c**) and corresponding model (**d**) of the basal body structure in the Δ*fliR* mutant. As in Δ*fliP*, FlhA density is absent, indicating that both FliP and FliR are required for recruitment or stabilization of FlhA.

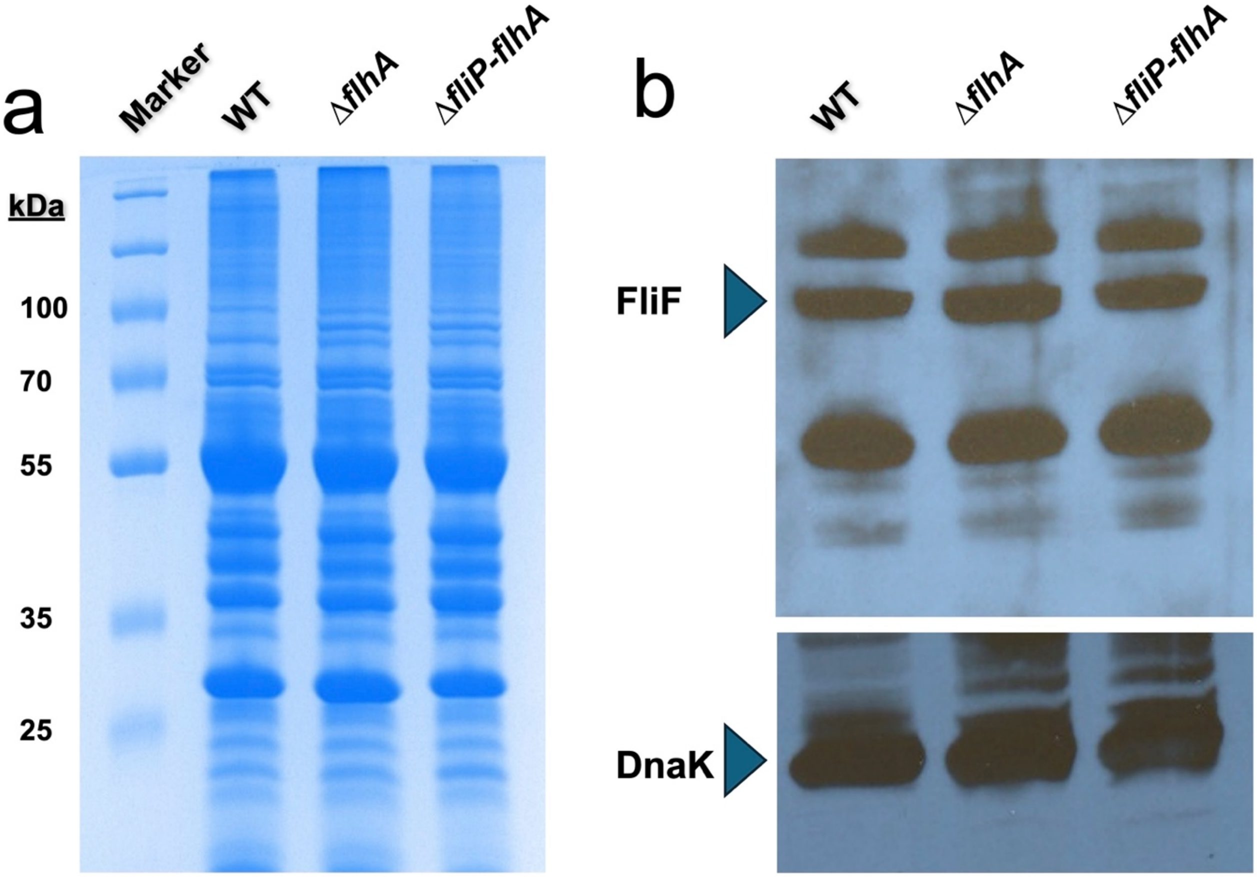


**Extended Data Fig. 7 | SDS–PAGE analysis.** (a, b) Equal amounts of *B. burgdorferi* cell lysates were analyzed by SDS–PAGE, followed by Coomassie blue staining or immunoblotting with specific antisera. Bands corresponding to FliF (65 kDa) and the loading control DnaK (72 kDa) are indicated.

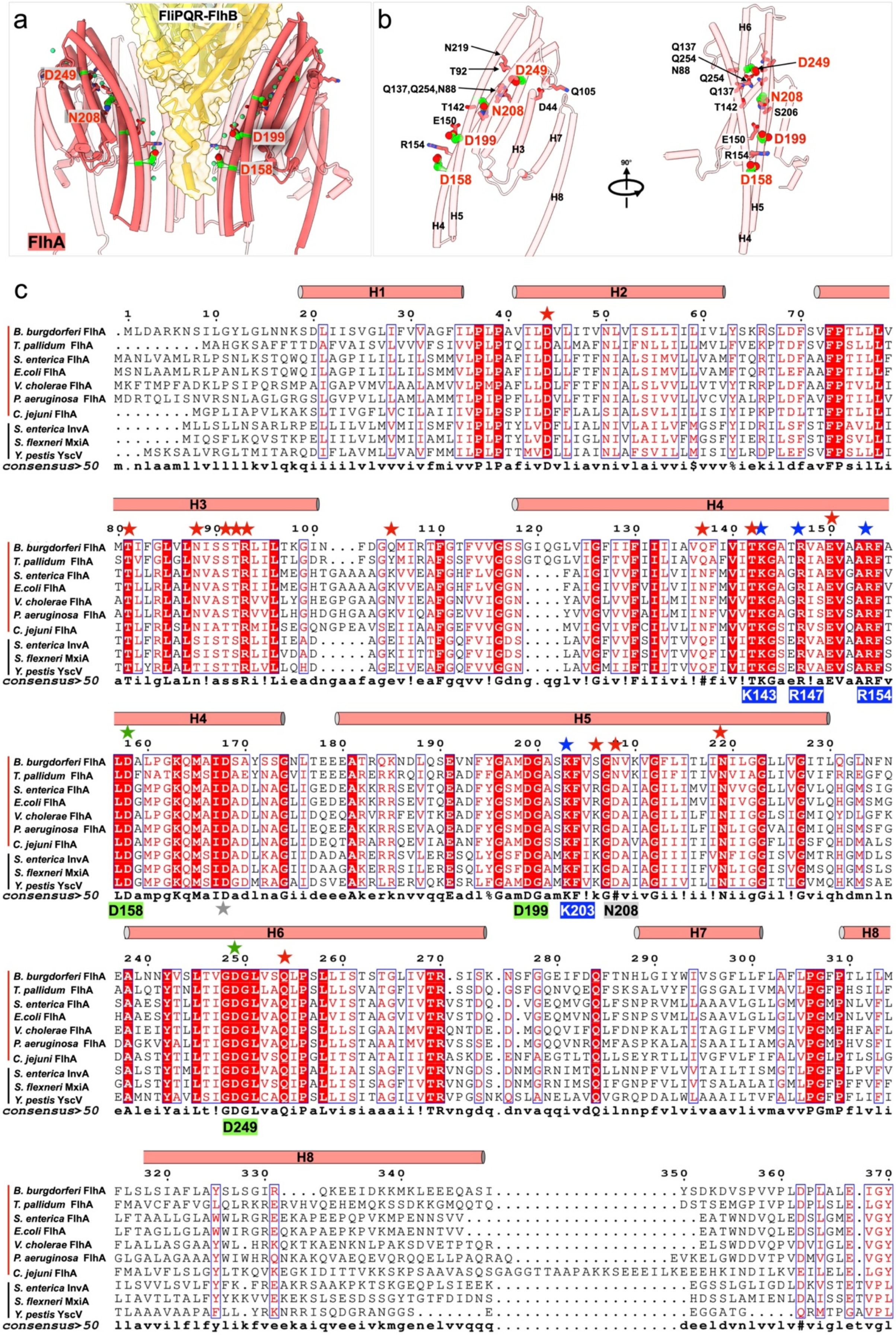
a
FliPQR-FlhB
D249
N208
D199
D158
FlhA
b
N219
T92
D249
Q137,Q254,N88
Q105
T142
N208
D44
E150
R154
D199
H3
H7
D158
H8
H5
H4
H6
Q137
Q254
N88
S206
90°
c
H1
H2
H3
H4
H5
H6
H7
H8
B. burgdorferi FlhA
T. pallidum FlhA
S. enterica FlhA
E.coli FlhA
V. cholerae FlhA
P. aeruginosa FlhA
C. jejuni FlhA
S. enterica InvA
S. flexneri MxiA
Y. pestis YscV
consensus>50
K143
R147
R154
D158
D199
K203
N208
D249

**Extended Data Fig. 8 | Hydrophilic pathway of FlhA and its conservation.** (a) Hydrophilic pathway within FlhA, with lining residues shown as sticks. Key acidic residues are highlighted (green). (b) Enlarged view of the hydrophilic pathway. (c) Sequence alignment of FlhA showing conservation. Pathway residues are marked (black stars), with basic (blue) and acidic (green) residues indicated.

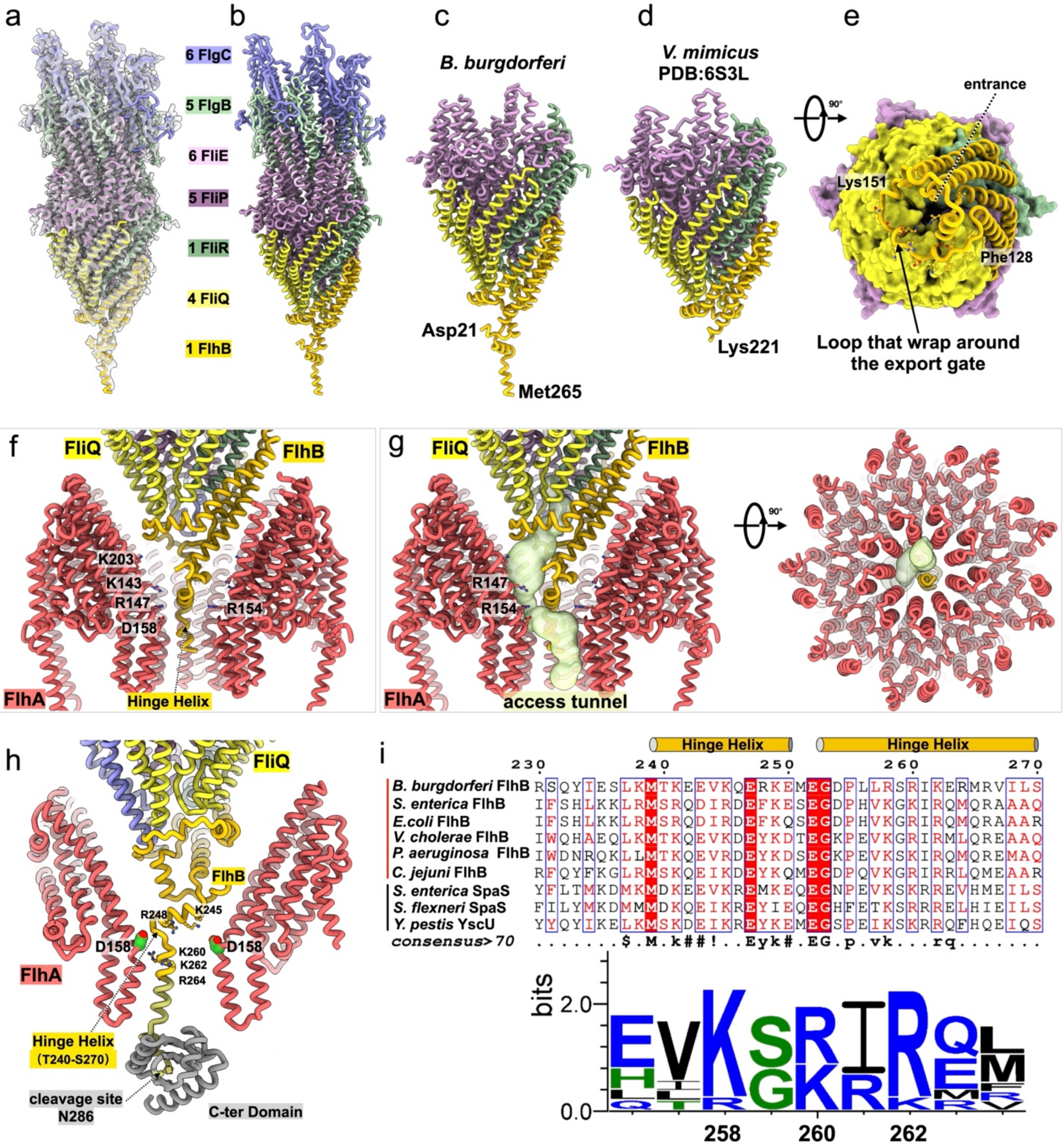


**Extended Data Fig. 9 | Spatial organization of the export gate and FlhA.** (a) Model of the export gate and proximal rod fitted into the cryo-EM map (transparent), with components and stoichiometries indicated. (b) Model of the export gate together with the proximal rod. (c, d) Structural comparison of the export gate from *B. burgdorferi* and *V. mimicus* (PDB: 6S3L). The *B. burgdorferi* structure contains an extended FlhB hinge helix absent from the *V. mimicus* model. (e) Enlarged view of an FlhB loop wrapping around the base of the export gate. (f) Spatial organization of FlhA relative to the FliPQR–FlhB export gate. Conserved basic residues surrounding the substrate entry site are indicated. (g) Access channel calculated using MOLEonline 2.0, illustrating the pathway through the FlhA tunnel to the export gate and its asymmetric orientation. (h) Full-length FlhB model based on the density map and an AlphaFold-predicted structure (grey), showing that the hinge helix is positioned toward one side of FlhA and enriched in basic residues. (i) Sequence alignment of the hinge-helix region highlighting enrichment of lysine and arginine residues.

**Extended Data Table 1| Bacterial strains used in this study**

| *B. burgdorferi* | Gene | Gene product | Motility | Reference |
|---|---|---|---|---|
| Wild type | N/A | N/A | motile | Zhao. *et al.* (2013) |
| Δ*fliP-flhA* | N/A | N/A | nonmotile | This study |
| Δ*flhA* | bb0271 | Flagellar biosynthesis protein FlhA | nonmotile | This study |
| Δ*flhB* | bb0272 | Flagellar biosynthesis protein FlhB | nonmotile | This study |
| Δ*fliP* | bb0275 | Flagellar biosynthesis protein FliP | nonmotile | This study |
| Δ*fliQ* | bb0274 | Flagellar biosynthesis protein FliQ | nonmotile | This study |
| Δ*fliR* | bb0273 | Flagellar biosynthesis protein FliR | nonmotile | This study |
| *flhA-D158N* | bb0271 | A point mutation in FlhA | nonmotile | This study |
| *flhA-D158E* | bb0271 | A point mutation in FlhA | less motile | This study |
| *flhA-D199N* | bb0271 | A point mutation in FlhA | motile | This study |
| *flhA-D249N* | bb0271 | A point mutation in FlhA | nonmotile | This study |

**Extended Data Table 2| Cryo-ET data collection and parameters**

| Genotype | No. of Tomograms | No. of Motors | Microscope/ Detector | Magnification | Pixel size (Å) | Resolution (Å) | Accession code |
|---|---|---|---|---|---|---|---|
| Wild type | 480 | 4322 | Titan Krios/ Gatan K3 | 42,000 | 2.15 | 15.9 | EMD-76383 |
| Δ*fliQ* | 735 | 4042 | Titan Krios/ Gatan K3 | 42,000 | 2.15 | 16.0 | EMD-48081 |
| Δ*flhA* | 429 | 3479 | Titan Krios/ Gatan K3 | 42,000 | 2.15 | 21.0 | EMD-48082 |
| *flhA-D158N* | 85 | 700 | Titan Krios/ Gatan K3 | 42,000 | 2.15 | N/A | N/A |
| *flhA-D158E* | 126 | 1009 | Titan Krios/ Gatan K3 | 42,000 | 2.15 | N/A | N/A |
| *flhA-D199N* | 101 | 541 | Titan Krios/ Gatan K3 | 42,000 | 2.15 | N/A | N/A |
| *flhA-D249N* | 68 | 409 | Titan Krios/ Gatan K3 | 42,000 | 2.15 | N/A | N/A |
| Δ*fliP-flhA* | 342 | 54 | Polara/CCD | 31,000 | 5.7 | 63.0 | EMD-6094 |
| Δ*flhB* | 543 | 217 | Polara/CCD | 31,000 | 5.7 | 58.0 | EMD-6089 |
| Δ*fliP* | 265 | 359 | Polara/CCD | 31,000 | 5.7 | 48.0 | EMD-6091 |
| Δ*fliR* | 229 | 830 | Polara/CCD | 31,000 | 5.7 | 41.0 | EMD-6093 |

**Extended Data Table 3| Cryo-EM Data Collection and Structure Refinement Statistics**

| | MS-ring (RBM3) | MS-ring (RBM1&2$_{inner}$) | MS-ring (RBM1&2$_{outer}$) | Export gate-rod |
|---|---|---|---|---|
| **Data collection** | | | | |
| Microscope | Titan Krios | Titan Krios | Titan Krios | Titan Krios |
| Voltage (kV) | 300 | 300 | 300 | 300 |
| Camera | K3 | K3 | K3 | K3 |
| Magnification | 81K | 81K | 81K | 81K |
| Total Electron Exposure ($e^-$/Å$^2$) | 70 | 70 | 70 | 70 |
| Defocus range (um) | -1.5 to -2.2 | -1.5 to -2.2 | -1.5 to -2.2 | -1.5 to -2.2 |
| Pixel size (Å) | 1.068 | 1.068 | 1.068 | 1.068 |
| Micrographs (no.) | 26,472 | 26,472 | 26,472 | 26,472 |
| **Data processing** | | | | |
| Symmetry imposed | C41 | C23 | C18 | C1 |
| Final particle images (No.) | 40,949 | 37,987 | 37,987 | 37,987 |
| Map resolution (Å) 0.143 FSC threshold | 3.37 | 3.25 | 6.42 | 3.87 |
| Map sharpening B factor (Å$^2$) | -97.1 | -87.7 | NA | -39.9 |
| **Refinement** | | | | |
| CC (model vs. data) | 0.73 | 0.78 | 0.79 | 0.82 |
| Non-hydrogen atoms | 86,264 | 37,536 | 29052 | 61,250 |
| Protein residues | 10,742 | 4,807 | 3,636 | 7,834 |
| Ligands | 0 | 0 | 0 | 0 |
| **B factors (Å$^2$)** | | | | |
| Proteins | 110.61 | 45.31 | 360.56 | 108.94 |
| **R.m.s. deviations** | | | | |
| Bond lengths (Å) | 0.004 | 0.004 | 0.002 | 0.003 |
| Bond angles (°) | 0.492 | 0.580 | 0.511 | 0.643 |
| **Validation** | | | | |
| MolProbity score | 1.97 | 2.27 | 1.64 | 2.02 |
| Clashscore | 5.10 | 6.24 | 4.32 | 4.25 |
| Rotamer outliers (%) | 3.06 | 4.91 | 3.56 | 3.76 |
| **Ramachandran plot** | | | | |
| Favoured (%) | 95.32 | 94.08 | 93.43 | 94.45 |
| Allowed (%) | 4.68 | 5.46 | 6.06 | 5.41 |
| Outliers (%) | 0.00 | 0.46 | 0.51 | 0.14 |
| PDB code | 12DR | 12DS | 12DT | 12DU |
| EMDB code | EMD-76344 | EMD-76345 | EMD-76346 | EMD-76347 |

| | FlhA$_{TMD}$ (1-350aa) | Overall model Export apparatus and MS-ring |
|---|---|---|
| **Data collection** | | |
| Microscope | Titan Krios | Titan Krios |
| Voltage (kV) | 300 | 300 |
| Camera | K3 | K3 |
| Magnification | 81K | 81K |
| Total Electron Exposure ($e^-$/Å$^2$) | 70 | 70 |
| Defocus range (um) | -1.5 to -2.2 | -1.5 to -2.2 |

| | | |
|---|---|---|
| Pixel size (Å) | 1.068 | 1.068 |
| Micrographs (no.) | 26,472 | 26,472 |
| **Data processing** | | |
| Symmetry imposed | C9 | / |
| Final particle images (No.) | 35,127 | / |
| Map resolution (Å) 0.143 FSC threshold | 4.39 | / |
| Map sharpening B factor (Å$^2$) | -165.9 | / |
| **Refinement** | | |
| CC (model vs. data) | 0.77 | / |
| Non-hydrogen atoms | 24,075 | 231,030 |
| Protein residues | 3,150 | 29,166 |
| Ligands | 0 | 0 |
| **B factors (Å$^2$)** | | |
| Proteins | 175.44 | / |
| **R.m.s. deviations** | | |
| Bond lengths (Å) | 0.002 | 0.002 |
| Bond angles (˚) | 0.503 | 0.499 |
| **Validation** | | |
| MolProbity score | 1.80 | 1.72 |
| Clashscore | 5.83 | 3.59 |
| Rotamer outliers (%) | 3.02 | 3.04 |
| **Ramachandran plot** | | |
| Favoured (%) | 97.41 | 96.72 |
| Allowed (%) | 2.59 | 3.14 |
| Outliers (%) | 0.00 | 0.14 |
| PDB code | 12DV | 12DW |
| EMDB code | EMD-76348 | EMD-76349 |
